# A large-scale analysis of the R2TP chaperone network reveals its contribution to the assembly of INO80, SRCAP and TIP60

**DOI:** 10.64898/2026.07.07.737026

**Authors:** Yoann Abel, Manon Philippe, Laurence Decourty, Ana C.F. Paiva, Philipp Busse, Marie-Cécile Robert, Serge Urbach, Camille Béllières, Franck Vandermoere, Jacques Imbert, Martial Seveno, Cosmin Saveanu, Pedro Sousa, Séverine Boulon, Tiago Bandeiras, Edouard Bertrand, Céline Verheggen

## Abstract

HSP90/R2TP is an essential quaternary chaperone composed of RPAP3, PIH1D1 and the RUVBL1/RUVBL2 AAA+ ATPases. These enzymes are also part of the INO80, SRCAP and TIP60 complexes, but the relationship between these chromatin remodelers and R2TP remains unclear. Here, we performed systematic analyses of the R2TP-specific subunits RPAP3 and PIH1D1. We validated 115 interaction partners and found that many were sensitive to HSP90 or R2TP inhibition. In yeast, epistatic screens revealed functional interactions with Ino80, Swr1 (SRCAP) and NuA4 (TIP60). Consistently, human RPAP3 physically interacted with subunits of INO80, SRCAP and TIP60 and was required for the formation of these complexes. More specifically, RPAP3 enabled the co-translational association of RUVBL1/RUVBL2 with the motor subunit of these chromatin remodelers. *In vitro*, the client-binding domain of RUVBL1/RUVBL2 modulated their interaction with RPAP3, suggesting that client subunits displace RPAP3 from nascent complexes. Thus, R2TP is an early chaperone of TIP60, SRCAP and INO80, which leaves RUVBL1/RUVBL2 as resident scaffolding subunits.

## Introduction

Protein chaperones such as HSP70 and HSP90 assist the folding of newly synthesized polypeptides in order to prevent their aggregation or degradation ^1^. Most of the time, binding to their clients is helped by co-chaperones that provide specificity and regulate the nucleotide hydrolysis cycle of the chaperone^2^. One of the key co-chaperones of HSP70 and HSP90 is R2TP, which performs a unique function in the cell by promoting the assembly of multi-subunit complexes. Notably, R2TP was shown to chaperone the assembly of the three nuclear RNA polymerases ^3,4^, snoRNPs ^5,6,7,8^, U4 and U5 snRNPs ^9,10,11^, the PIKK-containing complexes ^12,13^ and the TSC complex ^14^. Cells also contain R2TP-like chaperones exhibiting a subunit composition related to that of R2TP ^15^, and an important class of clients of R2TP and R2TP-like chaperones are the axonemal dyneins (reviewed in ^16,17^).

The R2TP consists of a RPAP3/PIH1D1 heterodimer associated with a hetero-hexameric ring of the AAA+ ATPases RUVBL1 and RUVBL2 ^18,19,20^. These closely related enzymes have a chaperone activity on their own. In turn, RPAP3 and PIH1D1 recruit clients, HSP70, HSP90 and also regulate RUVBL1/RUVBL2 ATPase activity. Although the R2TP chaperone was the subject of many studies, how it works at the molecular level is not entirely understood. RUVBL1 and RUVBL2 have an insertion domain called domain II (DII) that is believed to play key roles in their chaperone activity (reviewed in ^21^). The DII domains can adopt inward or outward conformations and the switch between them depends on the nucleotide bound to RUVBL1/RUVBL2 (ADP or ATP; ^20,17,22^). This switch also controls the interactome of RUVBL1/RUVBL2, some partners binding the ATP-state while others only bind in absence of ATP ^19,20^. It is believed that these properties of RUVBL1/RUVBL2 play a key role in controlling intermolecular interactions to ensure the proper assembly of client complexes. RUVBL1/RUVBL2 may recruit multiple client subunits, stabilize assembly intermediates and also remove early assembly factors to enable assembly to proceed ^7,8, 17,23^. Interestingly, a drug called CB-6644 was recently shown to block the ATPase activity of RUVBL1/RUVBL2 ^24,22^. This drug blocks the assembly of R2TP client complexes, confirming the key role of RUVBL1/RUVBL2 in this process^24,25,26^

The R2TP chaperone is well conserved across evolution. In metazoans, the R2TP core however forms a larger complex called the PAQosome, which contains prefoldin-like proteins, WDR92/DNAAF10 and POLR2E (reviewed in ^4,18^). RPAP3 and PIH1D1 are also conserved but have evolved differently in yeast and humans, leading to a different organization of R2TP ^27^. Human RPAP3 is enlarged compared to its yeast ortholog Tah1p. It has two TPR domains that enable its association with both HSP90 and HSP70 ^28^. Additionally, human RPAP3 contains a C-terminal domain that binds RUVBL1/RUVBL2 at the face of the hexameric ring opposite to DII ^15,19,20^. This anchors RPAP3 to the RUVBL1/RUVBL2 ring and allows some flexibility in the position of its other domains, enabling RPAP3 to fold back onto RUVBL1/RUVBL2 to reach their DII domains ^19,29^. It also allows RPAP3 to change the orientation of its HSP90-binding region, most likely to adapt its various clients ^29^. PIH1D1 is the fourth subunit of R2TP. It has a CS domain that binds a short peptide of RPAP3 located after its TPR2 and that is only present in the RPAP3 isoform 1 ^28^. PIH1D1 also has an N-terminal PIH domain that binds phosphorylated peptides with a DpSDD/E consensus motif ^30^.

The R2TP chaperone can recruit subunits of client complexes in various ways. First, the phospho-binding pocket of PIH1D1 can recruit some clients ^30^. Second, RPAP3 can also recruit clients. For instance, it can directly bind the Dicer cofactor TARBP2 via its TPR1, in a manner compatible with HSP90 binding ^31^. Finally, many clients are recruited via specific adaptor proteins. For instance, the PIKKs interact with R2TP via the TTT complex ^32,12^, while the core snoRNP proteins NOP58 and SNU13 are recruited by the R2TP cofactors NOPCHAP1, ZNHIT6 and the heterodimer NUFIP1/ZNHIT3 ^33,5,7^.

Interestingly, RUVBL1/RUVBL2 are also associated with several other complexes unrelated to R2TP, such as the INO80, SRCAP and TIP60 chromatin remodeling complexes. These complexes are involved in DNA repair and transcriptional regulation, and they have an ATPase motor protein that engages nucleosomes to displace them ^34,35,36^. In particular, SRCAP and TIP60 trigger the replacement of H2A by H2A.Z, an essential histone variant that is enriched at transcription start sites, while INO80 performs the opposite reaction (reviewed in ^37,38^). INO80 can additionally slide nucleosomes and place them adjacent to nucleosome-free regions, while TIP60 can acetylate H2A and H4 ^39,40^. These chromatin remodeling complexes play essential roles in the cell but their functional link with R2TP has remained unclear. In these complexes, RUVBL1/RUVBL2 are also found as hetero-hexameric rings as in R2TP. They are however devoid of RPAP3 and PIH1D1 and instead appear to provide a scaffolding role to these complexes ^41,42^. Overall, these activities are thought to represent independent functions of RUVBL1/RUVBL2.

Over the years, various proteomic screens have identified a number of R2TP putative partners, and thus potential additional clients and functions ^14,11,25^. In this study, we set out to refine R2TP function by performing an array of physical, genetic and functional assays. We validated 115 interaction partners of RPAP3 and PIH1D1 and found that many were sensitive to HSP90 inhibition. Most interestingly, we discovered genetic and physical interactions between RPAP3/PIH1D1 and INO80, and a detailed analysis showed that RPAP3 is required for the co-translational loading and subsequent association of RUVBL1/RUVBL2 with INO80, SRCAP and TIP60. Thus, these chromatin remodeling complexes are *bona fide* clients of R2TP, which keep RUVBL1/RUVBL2 as a resident chaperone.

## Results

### A yeast two-hybrid screen reveals ∼100 proteins interacting with RPAP3, covering a large range of functions

In order to uncover additional interactants of RPAP3, we first performed a yeast-two hybrid (Y2H) screen using the full-length RPAP3 protein as bait (Figure 1A). We screened a human placenta cDNA library and found 98 preys. A GO term enrichment analysis for their molecular functions or biological processes revealed a significant enrichment for chaperone activity, transcription, histone binding, RNA binding, nuclear import, cytoskeleton and protein phosphorylation (Figure 1B and S1B). The preys were then ranked from A (highest) to D (lowest) depending on their potential to be a real partner (Table S1, see methods). The ‘A’ proteins were ACTB, KIFAP3, PGD, VIM and WAC. RALBP1 was the only protein in the ‘B’ category, while CKAP2, TAF15, FHX and TRIM25 were ranked C. The majority of the proteins were in ‘D’ category. Nevertheless, this category contained well-characterized RPAP3 partners, such as TARBP2 from the RISC loading complex, which was shown to directly interact with RPAP3 ^31^. It also contained HSP90 whose C-terminal peptide directly binds RPAP3. In addition, the minimal interacting domains of these proteins, as defined from the Y2H data, nicely corresponded to the available structural information, highlighting the quality of the Y2H screen. In the case of HSP90, the screen revealed an additional RPAP3 interacting region in the M (middle) domain of HSP90, suggesting additional contact points besides the HSP90 C-terminal end. Interestingly, the ‘D’ category contained MORF4L2, a subunit of TIP60, and YY1AP1, a protein associated with YY1 in the INO80 complex. Several other ‘D’ proteins were involved in transcription (CDK12, CDK13, TAF6, MED7 and MED15), and DNA repair (RAD21 and WAPL from the cohesin complex, RAD50 from the MRE11/NSB1 complex, also known to bind PIH1D1 via a DpsDD/E motif; ^43^). Overall, these 98 proteins constitute potential new clients of the HSP90/R2TP system.

**Figure 1:**
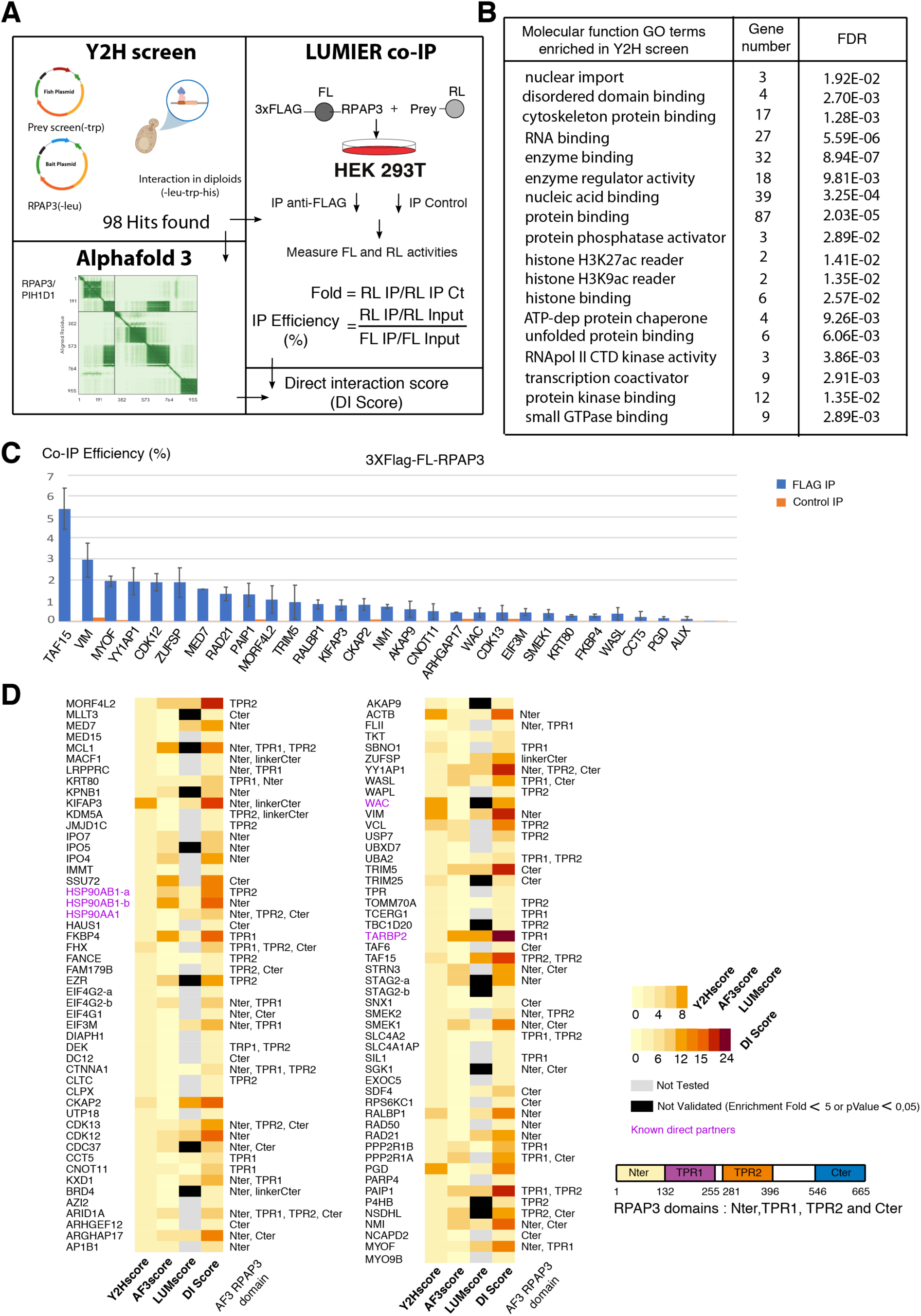
A yeast-two hybrid screen reveals a large number of RPAP3 interactants. **(A)** Schematic description of the assays used for classification of the preys found in Y2H screen using human RPAP3 as a bait. **(B)** GO term enrichment analysis for Molecular Functions for the RPAP3 interactants found in Y2H screen. The number of genes counted in each category and their false discovery rate (FDR) are indicated. **(C)** LUMIER co-IP assay showing the *in vivo* interaction between RPAP3 and a selection of the preys. The graph plots the co-IP efficiency of the indicated proteins when co-expressed with 3xFLAG-FL-RPAP3. The values of the blue bars are the IP/Input of the RL fusion proteins in the FLAG IP, normalized by the IP/Input values of the 3xFLAG-FL-RPAP3 fusion protein. The values of the orange bars are the IP/Input of the RL fusion proteins in the control IP, divided by the IP/input of the 3xFLAG-FL-RPAP3 fusion protein in the FLAG IP. Error bars: standard deviation (n=3). **(D)** Heat-map for all the preys found in the Y2H screen. For each protein, four different scores are given to evaluate the potential of the preys to make a direct physical interaction with RPAP3 (see methods). From left to right: Y2H score, AF3score and LUMscore (see Methods); Direct Interaction (DI) score is the sum of these scores. The RPAP3 domains with the smallest distance error with the prey, as predicted by AF3, are written on the right of the heat-map and a schematic of the RPAP3 protein is shown below the color bar. Some preys are present twice because two independent domains were found to interact with RPAP3 in the Y2H screen. Proteins in purple are previously known direct partners of RPAP3.

Next, we used additional methods to confirm the interactions found in the Y2H screen (Figure 1A). We first applied a structural analysis and screened each interaction using Alphafold 3 (AF3; Figure 1D). A score of 0 was given if no interaction was found or if the interacting region predicted by AF3 did not contain the domain found by Y2H. Otherwise, we assigned a score between 1 (lowest) and 8 (highest), depending on the minimal error predicted by AF3 for the distance between intermolecular interacting residues (see methods and Figure S1A). This gave a positive score for 38 proteins among the 98 Y2H hits (Figure 1D). Interestingly, AF3 also predicted which domain of RPAP3 bound the partner proteins and the N-terminal domain of RPAP3 was the most frequently involved (64% of the proteins tested). Next, we performed pairwise LUMIER co-immunoprecipitation (co-IP) assays. RPAP3 was fused at its N-terminus to a 3xFLAG tag and Firefly luciferase (FL), while the preys were fused with an HA tag and Renilla luciferase (RL). Among the 98 found in the Y2H screen, 56 ORFs were available in the ORFeome repository ^44^ and were tested in the LUMIER co-IP assay. Each prey plasmid was transiently co-transfected in HEK293T cells together with the vector expressing 3xFLAG-FL-RPAP3. Extracts were then immuno-purified with anti-FLAG antibodies, and FL and RL luciferase activities were measured in the immuno-purified and input fractions, in order to compute a co-IP efficiency: the IP/Input ratio of the prey normalized to that of the bait. We performed triplicate experiments and applied a stringent validation threshold of: (i) an enrichment of the prey in the FLAG IP higher than five times the control IP; (ii) a p-value lower than 0.05. Among the 56 preys tested, 21 had co-IP efficiencies higher than 0.32% (and up to 5.5%, Figure 1C). Finally, for the proteins tested, we computed a LUMscore based on the co-IP efficiency (Figure 1D, see methods). We then combined all three methods and computed a Direct Interaction score (DIscore) ranging from 2 to 24, with an equal number of points attributed for each method (Y2H, AF3, LUMIER co-IP; Figure 1D). Among the proteins with a score higher than 9, representing high confidence direct partners of RPAP3, we found TARBP2, a direct partner of R2TP ^31^, as well as proteins involved in transcription (TAF15,YY1AP1, MED7 and CDK12), chromatin folding (RAD21/cohesion) or cell architecture (VIM, CKAP2, MYOF).

### Systematic exploration of proteomic data by LUMIER co-IPs validates a large number of RPAP3 and PIH1D1 partners

The previous results suggested that RPAP3 may have more partners than previously anticipated. We thus aimed to validate the putative RPAP3 and PIH1D1 interactants found in previous proteomic analysis of R2TP ^7,10,11^. We generated a list of putative RPAP3 partners and collected a library of 172 plasmids containing the corresponding ORFs (different from the 56 ORFs from the Y2H screen). These ORFs were also fused with HA tag and Renilla luciferase (RL) and tested by LUMIER co-IP with 3xFLAG-FL-RPAP3 (Figure 2A). We found a total of 115 proteins that were considered as true interactants of RPAP3, i.e. with an enrichment over the control IP higher than 5 and a corresponding p-value lower than 0.05 (Figure 2B, Table S2; see methods). Of these, 74 proteins had a co-IP efficiency higher than 0.5% (Figure 2C and S2A). Among the proteins with the highest LUMIER co-IP efficiency, we found members of the PAQosome (URI1, 7.64%; UXT, 2.11%; and WDR92 8.86%), and known R2TP clients such as the U5 protein EFTUD2 (1.82% of co-IP efficiency), the core snoRNP protein NOP58 (0.78%) and the largest subunit of RNA polymerase II (POLR2A, 1.04%). Interestingly, we also found proteins not previously described as RPAP3 partners, such as the paraspeckle proteins NONO (7.22%) and PSPC1 (2.32%), several RNA polymerase II associated factors (DDX21, 4.87%; TAF15, 6.45%; CDK12, 1.87%; and MED7, 1.22%), the stress granule forming protein G3BP1 (3.07%) and, interestingly, the chromatin remodeler INO80 (5.92%).

**Figure 2:**
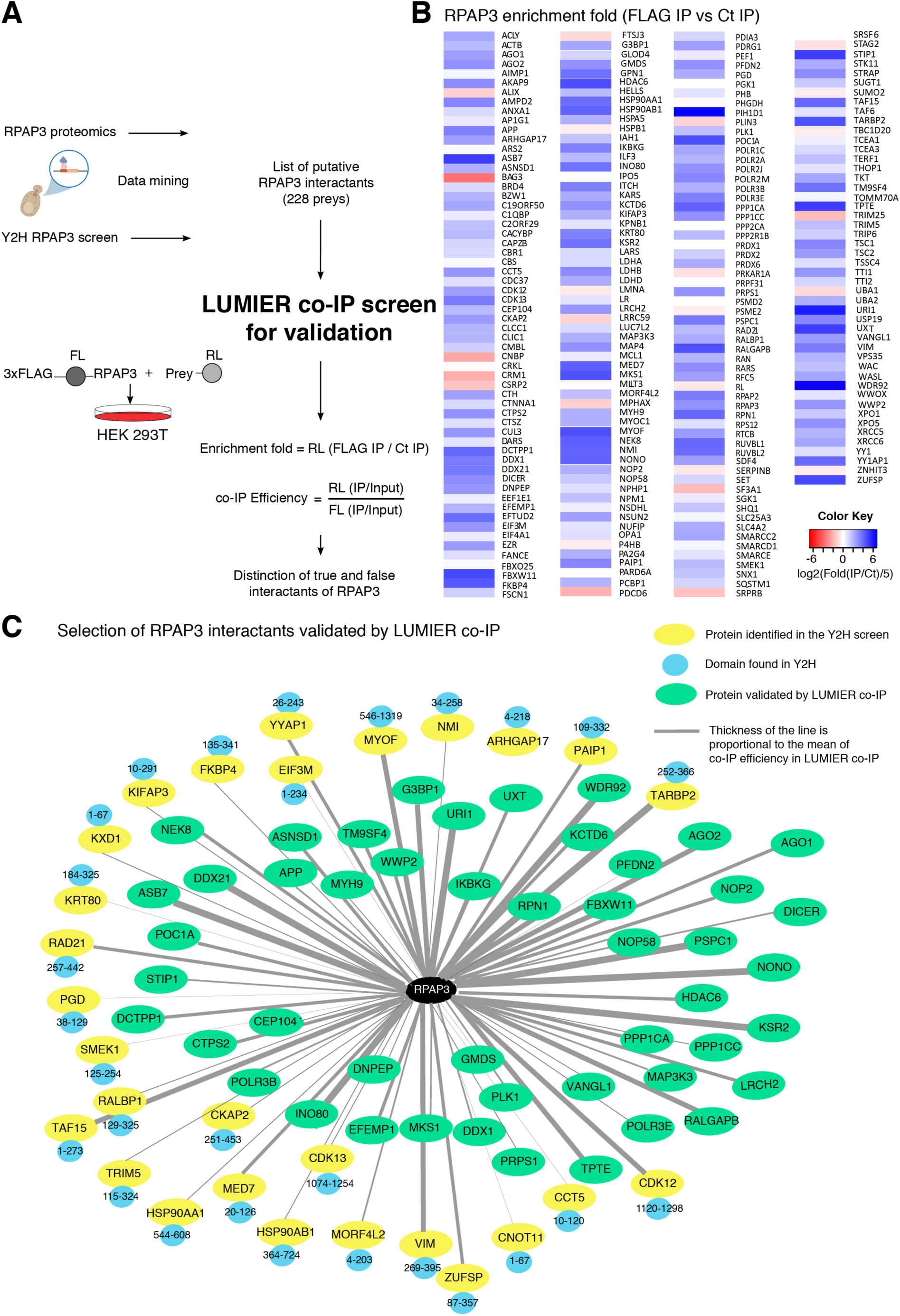
A medium-throughput LUMIER co-IP screen validates many RPAP3 partners. **(A)** Schematic representation of the LUMIER co-IP assay highlighting the choice of the preys and the metrics measured. True interactants have a enrichment fold >5 associated with a pValue < 0.05 (n=3). **(B)** Heat map for all preys tested by LUMIER co-IP with 3xFLAG-FL-RPAP3. The enrichment fold (RL in the FLAG IP versus the control IP) is represented for each protein tested (shown as Log2(enrichment fold/5); n=3). **(C)** Graphical representation of the proteins found to interact most strongly with 3xFLAG-FL-RPAP3 by LUMIER co-IP (Enrichment fold >5, % of co-IP efficiency >0.5, pvalue <0,05). The thickness of the grey lines represents the % of the co-efficiency of the interaction tested (from 0.5 to 7.5% for all the proteins represented). Proteins that were first identified in the Y2H screen with RPAP3 as a bait were shown in yellow bubbles and for each protein, the fragment found in Y2H assay is also indicated in an attached blue bubble. Proteins previously found in the interactome of RPAP3 by proteomic analysis and not in Y2H screen are shown in green bubbles.

Next, we performed a similar analysis for PIH1D1 partners, using available PIH1D1 proteomic data ^10,14^. We tested 186 preys for their interaction with PIH1D1 in LUMIER co-IPs, this time using a single assay (Figure 3A, Table S2). Due to the absence of replicate, we considered as validated interactants the proteins with an enrichment in the FLAG IP versus the control IP higher than 15 (Figure 3B). These validated 49 PIH1D1 partners, of which 26 had a co-IP efficiency higher than 0.5% (Figure 3C and S3A). These factors included PAQosome proteins (WDR92), known R2TP co-factors or PIH1D1 partners (ECD, RPAP2, GNP3; TSC1, TSC2; ^3,14,30^), partners also found in the RPAP3 LUMIER experiments (POLR2A, POLR3E, NONO, PSCP1), as well as a series of 14 novel PIH1D1 partners. Interestingly, these included the RISC proteins AGO1 and AGO2, suggesting that these interactions may be related to the role of R2TP in miRNA metabolism ^31^.

**Figure 3:**
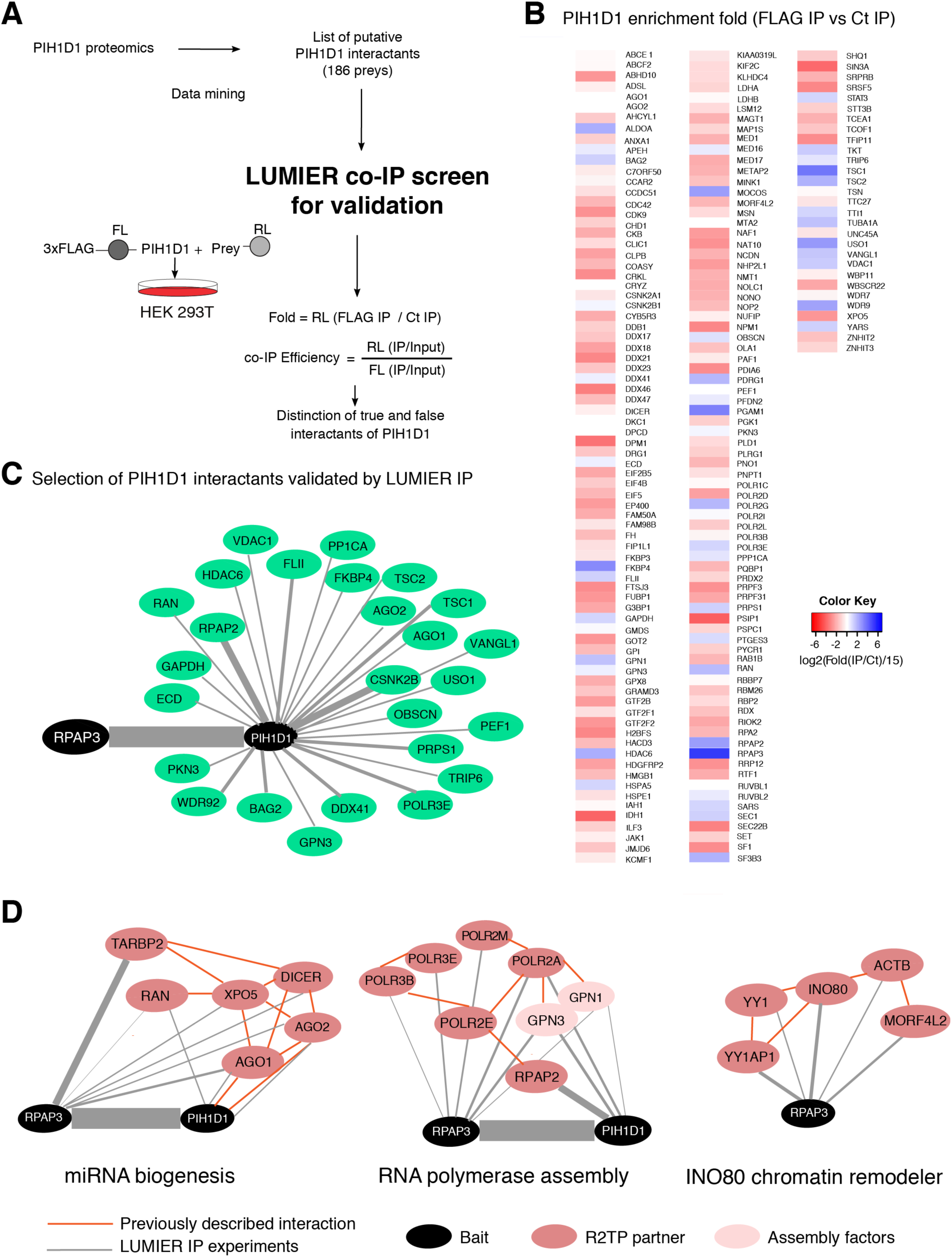
A medium-throughput LUMIER co-IP screen validates PIH1D1 partners. **(A)** Schematic representation of the LUMIER co-IP assay highlighting the choice of the preys and the metrics measured. True interactants have a Fold >15, n=1. **(B)** Heat map for all preys tested by LUMIER co-IP with 3xFLAG-FL-PIH1D1. The enrichment fold (RL in the FLAG IP versus the control IP) is represented for each protein tested (shown as Log2(enrichment fold/15); n=1). **(C)** Graphical representation of the proteins found to interact most strongly with 3xFLAG-FL-PIH1D1 by LUMIER co-IP. The thickness of the grey lines represents the % of the co-efficiency of the interaction tested. **(D)** Selected network of proteins interacting with RPAP3 and PIH1D1, and involved in miRNA biogenesis and function (left), RNA polymerase assembly (middle) and the INO80 complex (right). The preys shown have a LUMIER co-IP efficiency > 0.5% and an enrichment fold over the control IP > 5 for RPAP3 and >15 for PIH1D1. Red line: interactions documented in String or in the literature.

In order to identify a comprehensive set of client complexes potentially assembled by R2TP, we performed a GO term enrichment analysis for the ‘Cellular Component’ category, using all the newly validated partners of RPAP3 and PIH1D1 (Figure S2B). This revealed expected GO terms, such as the R2TP/prefoldin complex, the TTT complex, RISC, the RNA polymerases and the TSC complex, axonemal dyneins, as well as less expected ones, such as the Ku70/Ku80/DNAPK complex, Cyclin/CDK transcription elongation complex, PTW/PP1 phosphatase complex, tRNA splicing ligase complex, paraspeckles, and also a series of chromatin remodelers such as B-WICH, INO80 and the TIP60 chromatin remodeling complex (e.g. NuA4). This prompted us to explore in greater depth the interaction networks of selected categories, related to miRNA biogenesis, RNA polymerase assembly, U5 snRNP assembly, INO80, and paraspeckles (Figure 3D and S2C). This revealed a dense interaction network linking R2TP with proteins involved in miRNA maturation and function: Ran, XPO5, DICER, TARBP2, AGO1 and AGO2. Likewise, factors involved in the assembly of RNA polymerases showed a rich interaction network with multiple subunits and assembly factors connected to RPAP3 and PIH1D1. Interestingly, we also found multiple proteins of the INO80 chromatin remodeler complex connected to RPAP3 (INO80, MORF4L2, YYAP1 and ACTB), suggesting that RPAP3, and not only RUVBL1/RUVBL2, might somehow be linked to this complex. Together, these data indicate that the targets of R2TP extend beyond the currently known clients, and that its function in assembly likely relies on multiple interactions with subunits and assembly factors of its client complexes.

### Inactivation of HSP90 and R2TP affects the levels of RPAP3 and PIH1D1 interactants

To test whether the newly validated partners of RPAP3 and PIH1D1 were functionally relevant, we analyzed their fate after inactivation of HSP90 and R2TP. We used several conditions (Figure 4A): (i) treatment with Geldanamycin to inhibit HSP90; (ii) PIH1D1 KO cells ^14^; (iii) cells depleted for RPAP3 using an auxin-inducible degron system (RPAP3-mAID; ^31^). We selected 114 proteins among the validated interactants of RPAP3 and PIH1D1 plus few already known partners and used the fusion of their ORFs to HA-RL to easily measure their expression levels. The fusion proteins were transiently expressed from a moderate L30 promoter, and co-transfected with a Firefly luciferase (FL) expression vector to normalize for transfection efficiency. RL/FL ratios were then measured in cellular extracts done in triplicates and the fold-change of expression from the treated versus the control condition was further normalized using a non-relevant control protein (mPHAX; Figure 4A). For Geldanamycin treatment and PIH1D1 KO cells, non-treated parental cells were used as controls. In contrast, for the RPAP3-mAID experiment, we used wild-type matched HCT116 cells as controls, as untreated RPAP3-mAID cells exhibit reduced RPAP3 levels compared with wild-type cells even in the absence of auxin ^31^, and may therefore be hypomorphic for R2TP activity. Overall, only 29 out of the 114 proteins were not affected by any of the treatments (Figure 4B and S3B, Table S3). We found that 53 proteins were significantly less expressed in presence of Geldanamycin, whereas only 6 were more expressed, consistent with the previously proposed role of HSP90 in stabilizing unassembled R2TP clients ^5^. A similar pattern was observed for the depletion of RPAP3, with 33 proteins less expressed in its absence (while 12 were more expressed). Interestingly, the pattern was opposite with PIH1D1, with only 7 proteins less expressed in the KO cell line, while 38 were more expressed. This may suggest a role of PIH1D1 in the degradation of R2TP clients, possibly for the purpose of quality control. In conclusion, we found that many of the RPAP3 and PIH1D1 partners were sensitive to HSP90 inhibition and were thus putative clients of this chaperone. R2TP inactivation had less effects, which differed depending on the subunit that was removed.

**Figure 4:**
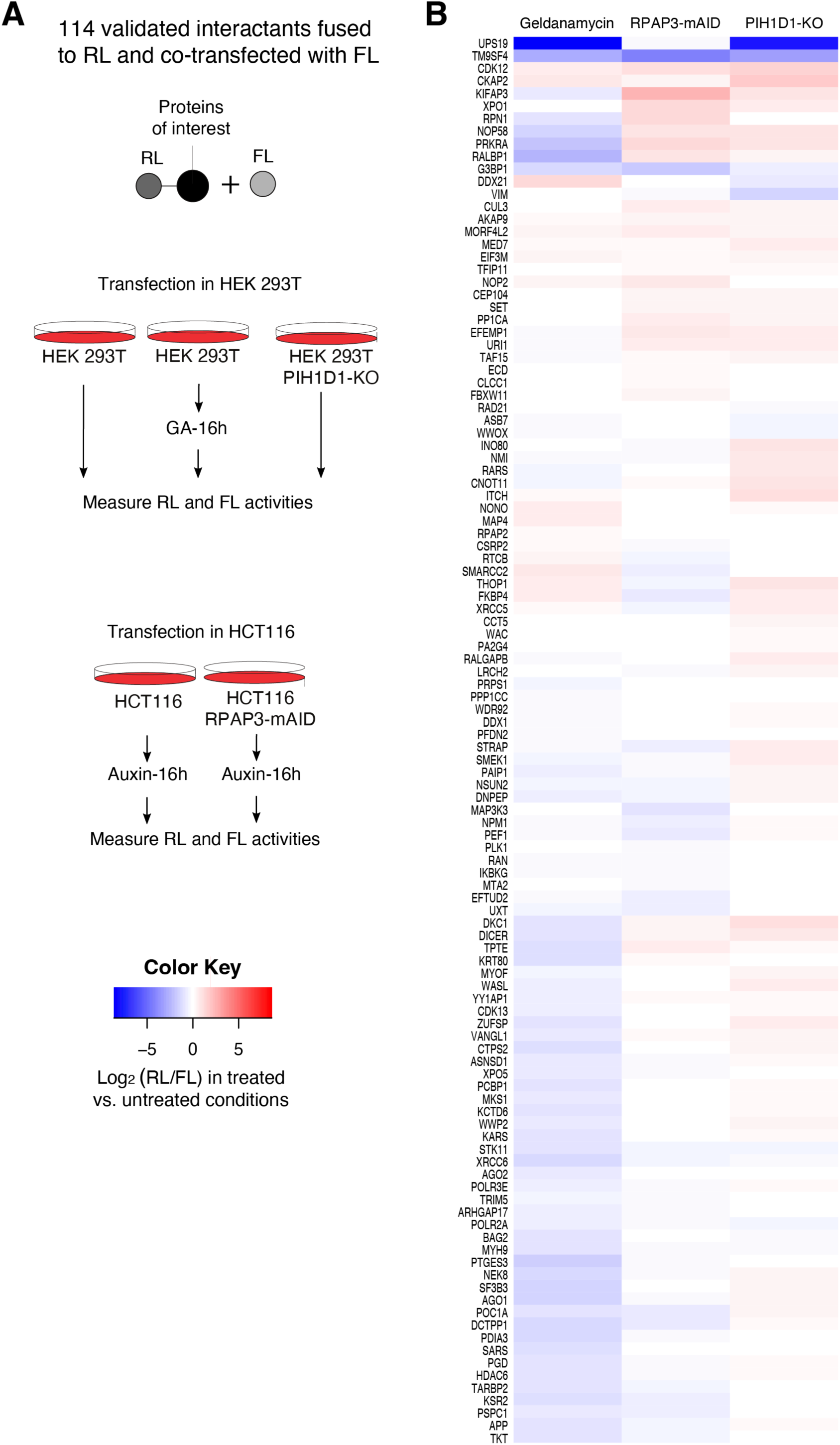
Effect of R2TP inhibition on the expression of RPAP3 and PIH1D1 partners. **(A)** Schematic representation of a functional expression screen done with 114 partners of RPAP3 and PIH1D1. Plasmids expressing the different proteins in fusion with RL were co-transfected with an FL expression plasmid to normalize for transfection efficiencies. HSP90/R2TP was inhibited in three different ways: HSP90 was inhibited by treating cells with Geldanamycin (GA) for 16h, R2TP was inhibited by genetically knocking-out of PIH1D1 (PIH1D1 KO cells) or by treating RPAP3-mAID degron cells with a derivative of auxin for 16h. RL level for mPHAX-RL fusion was measured in each condition and used for normalization. **(B)** Heat-map showing the effects of R2TP inhibition on expression of RPAP3 and PIH1D1 partners. The values are coded according to the color bar shown in A, and represent the RL/FL activity ratio in the treated conditions divided by the one in the control condition, further divided by the value obtained for mPHAX to normalize for difference in cell lines. The values are averages of three experiments and are given as Log_2_.

### Genetic interaction mapping of the yeast R2TP subunit Pih1 reveals links with the Ino80, Swr1 et NuA4 chromatin remodeling complexes

To better identify the function of RPAP3 and PIH1D1, and in particular to extend our analyses beyond the expression levels of their partners, we turned to yeast to perform a genetic interaction screen using a PIH1 deletion strain, *S. cerevisiae* homolog of PIH1D1. The deletion strain was mated with a pool of mutants of the yeast systematic deletion library of non-essential genes and with a combination of DAmP mutants for essential genes^45,46,47^.

The PIH1 deletion led to a range of strong negative and positive genetic interaction results that were well correlated among two independent replicate experiments (Figure 5A). We observed a strong negative genetic interaction between *pih1*Δ, and a hypomorphic allele of RVB1 (*rvb1*-DAmP), the ortholog of RUVBL1, validating the approach. We also found negative genetic links of *pih1*Δ with DAmP allelles of NOP58, NHP2 and CBF5, which code for core snoRNP proteins (Figure 5A-B and Table S4). These negative interactions nicely fit the known role of R2TP in snoRNP assembly ^5,6^. Epistatic or suppressor interactions also included deletion for genes coding for ribosomal proteins, RNA metabolism genes (e.g. the LSM complex), or the COMPASS histone methylation complex.

**Figure 5:**
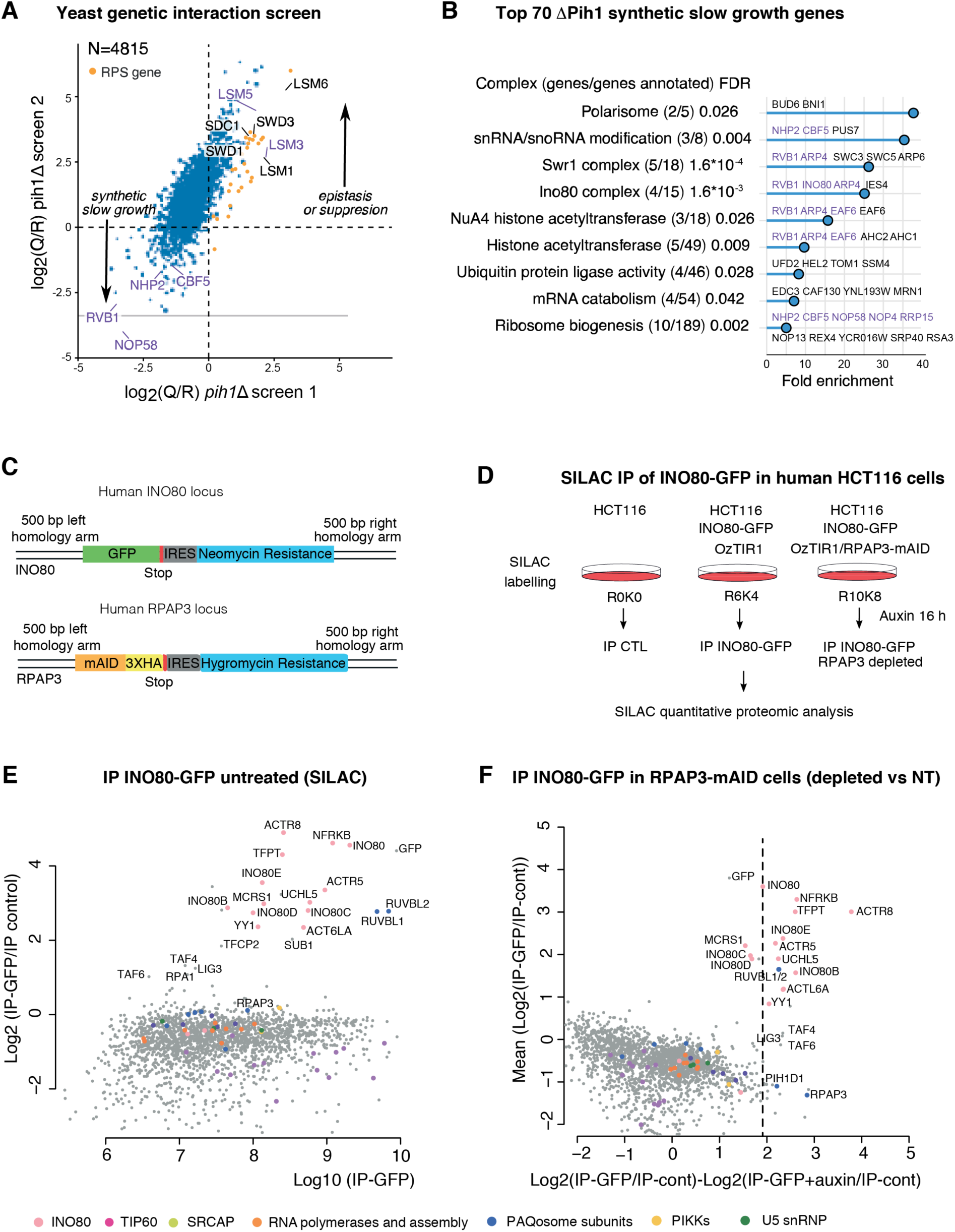
Genetic interaction mapping with deletion of PIH1 in yeast and impact of RPAP3 depletion on the assembly of INO80 complex in HCT116 cells. **(A)** Results of two independent genetic interaction screens using the deletion of PIH1 in *S. cerevisiae* combined with the systematic deletion and DAmP modification of genes. Axis values represent normalized and pleiotropy corrected log_2_(Q/R), where Q represents the signal intensities of the tag marking a given mutant when combined with the query mutation, and R is the signal from the same mutant when introduced in a reference population. Negative values indicate a synthetic slow growth defect, while positive values indicate suppression of a slow growing phenotype or epistasis (most frequent). Orange dots correspond to genes of ribosomal small subunit proteins. Several genes of interest are annotated in black for deletion mutants and in purple for DAmP alleles. **(B)** GO terms (cellular component) enriched in the 70 top genes that showed a negative genetic interaction with the deletion of PIH1 and a log_2_(Q/R) inferior to -1 in both screens. For each term, the number of genes from the list matching with the term and the total number of genes annotated to that term are indicated, together with the fold enrichment and the corresponding false discovery rate (FDR), as reported by ShinyGO ^48^. Purple gene names indicate DAmP modification. **(C)** Schematic of the cassettes integrated at the INO80 and RPAP3 genomic loci. The RPAP3-mAID is homozygous while INO80-GFP is heterozygous. **(D)** Schematic of the experiment for the SILAC proteomic analysis of the INO80-GFP IP in RPAP3-mAID depleted cells vs parental cells. **(E)** The graph represents the enrichment in the INO80-GFP IP versus the control IP (y axis; Log2) as a function of signal intensity (x axis; Log10), as measured in the parental HCT116 cells expressing INO80-GFP. Each dot represents a protein that is colored according to the code below the graphs. **(F)** The graph shows the enrichment of proteins in the INO80-GFP IP versus the control IP (y axis, mean of auxin-treated parental and RPAP3-mAID cells), as a function of their differential enrichment in the RPAP3-mAID depleted vs the parental cells, for the same IPs (x axis, Log2 (ratio)).

To identify the macromolecular complexes affected by the deletion of PIH1, we performed a GO term enrichment analysis for the category ‘Cellular Component’ ^48^, using the 70 gene deletions or DAmP strains most negatively affected by PIH1 deletion. Surprisingly, this analysis identified three chromatin remodeling complexes, SWR1 (the ortholog of SRCAP, enrichment FDR 1.6*10^-4^), INO80 (FDR 1.6*10^-3^) and NuA4 (FDR 0.026; Figure 5B). NuA4 acetylates histone H4 and is related to the human TIP60 complex, with the notable difference that NuA4 does not contain the Rvb1/Rvb2 AAA+ ATPases^49^. Additionally, a systematic analysis confirmed that many subunits of these complexes had negative genetic interactions with PIH1, like HTZ1, which codes for the histone variant H2A.Z, a key substrate of INO80 and SWR1 (Figure S4A). These data, together with the physical interaction of INO80 with RPAP3 that we observed in human cells (Figure 2B), raised the possibility that R2TP may be directly involved in the biogenesis of INO80, SWR1/SRCAP and NuA4/TIP60.

### RPAP3 is required for the assembly of the INO80, SRCAP and TIP60 complexes

To gain more insight into the role of R2TP in the biogenesis of the INO80 complex, we analyzed the effect of RPAP3 depletion on INO80 integrity. We used the HCT116 cell line containing an homozygous RPAP3-mAID allele and the parental HCT116 cells ^31^, and further modified these cells by inserting by CRISPR a GFP tag at the C-terminus of the endogenous INO80 subunit (Figure 5C and S4B). We then IP’ed INO80-GFP and compared its partners in presence and absence of RPAP3 by quantitative mass spectrometry (Figure 5D). In the parental HCT116 cells, all the subunits of the INO80 complex were detected in the INO80-GFP IP, with a 4-20 fold enrichment over the control IP (Figure 5E, Table S5). However, when RPAP3 was removed by treating the INO80-GFP RPAP3-mAID degron cells with a derivative of auxin for 16h, many subunits of the INO80 complex became less associated with INO80-GFP. This effect of RPAP3 removal is particularly notable for the key subunit ACTR8 of the HSA INO80 module, for NRFKB and TFPT that are part of the N-ter INO80 module, and for INO80B, part of the C module ^50^.

To confirm these results, we repeated the same experiment but using RUVBL1 as a bait instead of INO80. We used our previously generated HCT116 cells having a GFP tag fused to the endogenous C-terminus of RUVBL1 ^25^, and further modified this cell line to introduce a dTAG-3xHA at the C-terminus of RPAP3 alleles (Figure 6A and S5A). Proper homozygous insertion of the dTAG was verified by PCR genotyping and Western blots showed that RPAP3 was strongly depleted after treatment with dTAG-V1 (Figure S5B). In addition, the basal levels of RPAP3-dTAG were much less affected than with the RPAP3-mAID degron, indicating that this cell line is a better tool to study the effects of the rapid depletion of RPAP3 (Figure S5B and ^31^). We then performed label-free AP-MS of RUVBL1-GFP with and without a dTAG-V1 treatment for 2h (Figure 6C). In untreated conditions, the enrichment value of the RUVBL1-GFP IP versus a control IP showed the expected partners (Figure S5C, Table S6; compare with Figure 2A of ^25^: (i) R2TP clients, with the subunits of the nuclear RNA polymerases, U5 snRNP, snoRNPs and their assembly factors; (ii) R2TP/PAQosome subunits (RPAP3, PIH1D1 and the WDR92/prefoldin module; (iii) subunits of the INO80, SRCAP and TIP60 chromatin remodeling complexes. Importantly, short treatment with dTAG-V1 led to a ∼30-fold loss of RPAP3 from RUVBL1-GFP, concomitant with a 30-60-fold loss of PIH1D1 and the prefoldin module of the PAQosome (Figure 6C, Table S6). The subunits of the polymerases and their assembly factors were the next group of the most affected proteins, with losses in the range of 4-10-fold, except for POLR2B that was not affected. Remarkably, many subunits of INO80, SCRAP and TIP60 were also affected by RPAP3 removal, with their association with RUVBL1-GFP decreasing from 2 to 4-fold. Indeed, a systematic analysis of all the subunits of these complexes indicated that most of them were lost from RUVBL1-GFP (Figure 6C). These results indicate that the removal of RPAP3 affects the integrity of these chromatin remodeling complexes.

**Figure 6.**
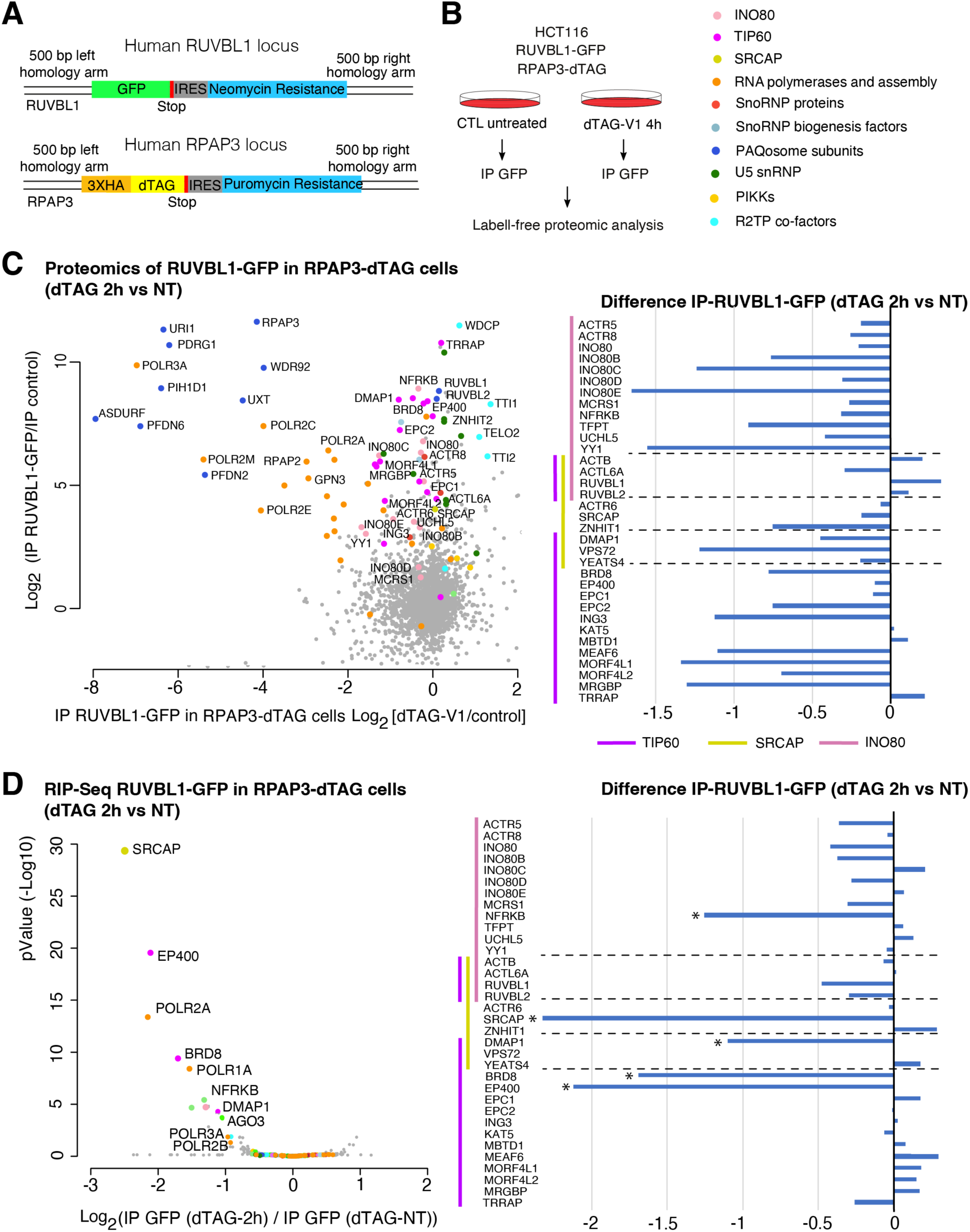
The rapid depletion of RPAP3 prevents the association of RUVBL1-GFP with its clients. **(A)** Schematic of the cassettes integrated at the RUVBL1 and RPAP3 genomic loci. The RPAP3-dTAG is homozygous and the RUVBL1-GFP is heterozygous. **(B)** Schematic of the experiment for the label-free proteomic analysis of RUVBL1-GFP IPs in presence and absence of RPAP3. **(C)** The graph shows the enrichment of proteins in the RUVBL1-GFP IP versus the control IP (y axis, mean of dTAG-treated and untreated cells), as a function of their differential abundance in the RPAP3-dTAG depleted vs untreated cells, for the same IPs (x axis, Log2 (ratio)). Each dot represents a protein that is colored according to the code on the right of panel B. The barplot shows the differential association of RUVBL1-GFP with all the subunits of the INO80, TIP60 and SRCAP complexes, with and without RPAP3 depletion. **(D)** The volcano plot shows the association of RUVBL1-GFP with various mRNAs in RPAP3-depleted versus non-depleted cells. The y axis shows the p-value (y axis; -Log10) for the detected mRNAs, as a function of their fold change in the RUVBL1-GFP IPs between dTAG-treated versus non-treated cells (x axis; Log2). Each dot represents a mRNA that is colored according to the code on the right of panel B. The barplot shows the differential association of RUVBL1-GFP with the mRNAs coding for all the subunits of the INO80, TIP60 and SRCAP complexes, in RPAP3-depleted versus non-depleted cells. *: p-values lower than 0.05.

### RPAP3 is important for the co-translational binding of RUVBL1/RUVBL2 to the large subunits of the INO80, SRCAP and TIP60 chromatin remodeling complexes

We previously showed that R2TP binds many of its client proteins while they are being translated ^25^. In addition, we observed that RUVBL1 associated co-translationally to several key subunits of the INO80, TIP60 and SRCAP chromatin remodeling complexes ^25^: EP400 and BDR8 of TIP60, the SRCAP motor subunit of SRCAP, and, to a lesser extent, the INO80 motor subunit of INO80. To study the role of RPAP3 in this process, we used the RUVBL1-GFP / RPAP3-dTAG cells to IP the endogenous RUVBL1-GFP and sequence the associated RNA, with and without addition of dTAG-V1 (RIP-Seq, Figure 6D and S5D, Table S7). Remarkably, depleting RPAP3 for 2h led to a strong decrease in the association of RUVBL1-GFP with many client mRNAs, including and foremost SRCAP, EP400, BRD8 mRNAs (4-8 fold decrease), and to a lower extent DMAP1 and NFRKB mRNAs (Figure 6D). The loss of binding also affected other families of R2TP clients, including the mRNAs coding for large RNA polymerase subunits and for the AGO1/AGO3 proteins. These results show that RPAP3 plays an important role in driving the co-translational loading of RUVBL1 on its clients, and this is especially important for the key subunits of the SRCAP and TIP60 chromatin remodelers. It is also interesting to note that the subunits affected by the removal of RPAP3 are not the same in the RIP-seq and proteomic experiments. In the RIP-seq, the large subunits are lost, while in the proteomic experiments, RUVBL1 remains largely associated with these subunits but loses other ones. This suggests that in absence of RPAP3, RUVBL1 can still bind the large subunits of SRCAP, TIP60 and INO80, but it does so post-translationally and this disrupts the formation of the full complexes.

### Formation of the INO80, SRCAP and TIP60 complexes require the ATPase activity of RUVBL1/RUVBL2

We previously showed that the nucleotide cycle of RUVBL1/RUVBL2 controlled the loading and release of these proteins on their clients ^25^. In particular, the effect of blocking the ATPase activity of RUVBL1/RUVBL2 prevented their co-translational loading. To better understand the role of the ATPase cycle of RUVBL1/RUVBL2 on the biogenesis of the INO80, SRCAP and TIP60 chromatin remodeling complexes, we reanalyzed our previous RUVBL1-GFP AP-MS experiments performed in presence of CB-6644 (^25^; Figure 7A), which is a specific inhibitor of the RUVBL1/RUVBL2 ATPase activity ^24^. We systematically measured the effect of CB-6644 on the association of RUVBL1-GFP with the subunits of INO80, SRCAP and TIP60 (Figure 7B). Remarkably, the addition of CB-6644 induced the dissociation of many subunits of these complexes from RUVBL1-GFP (28 subunits out of 34 detected). This was the case for all three complexes, with losses of up to 15-30-fold for INO80, INO80E, ACTR8, UCHL5 and TFPT.

**Figure 7.**
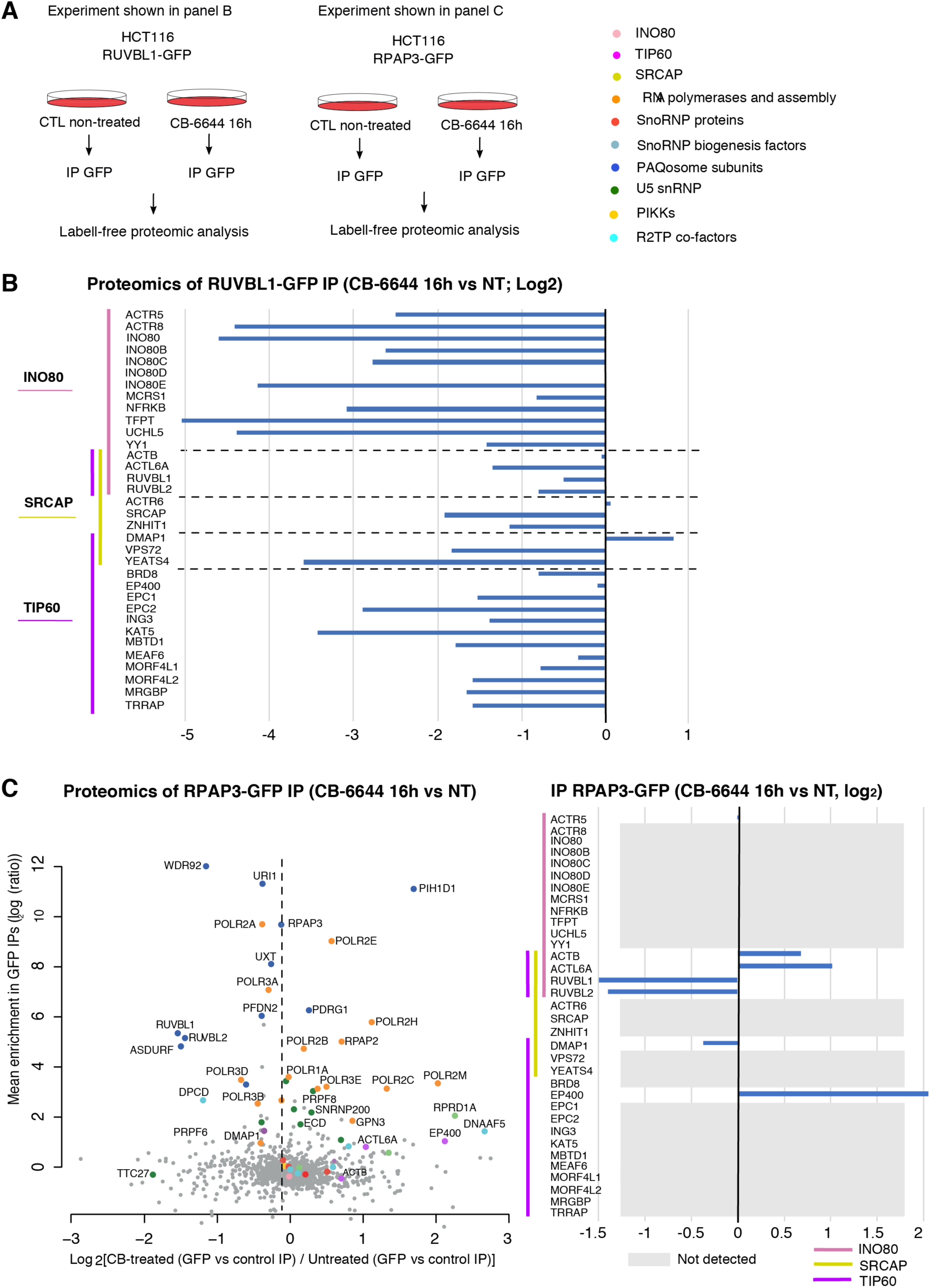
Impact of CB-6644 treatment on the association of RUVBL1-GFP and RPAP-GFP with their partners. **(A)** Schematic of the proteomic experiments of RUVBL1-GFP and RPAP3-GFP partners by label-free proteomics in HCT116 cells, treated or not with CB-6644 for 16h. **(B)** Barplot representing the fold change of the association of RUVBL1-GFP with all subunits of the INO80, SRCAP and TIP60 complexes, in CB-6644 treated versus untreated cells. The fold changes were measured by label-free proteomic analysis of RUVBL1-GFP IPs. **(C)** The graph shows the enrichment of proteins in the RPAP3-GFP IP versus the control IPs (y axis, mean of treated and untreated conditions), as a function of their differential abundance in the CB-6644 treated versus untreated cells, for the same IPs (x axis, log2 (ratio)). Each dot represents a protein that is colored according to the legend shown in A. The barplot shows the fold change of the association of RPAP3-GFP with all subunits of the INO80, TIP60 and SRCAP complexes, in CB-6644 versus untreated cells. The fold changes were measured by label-free proteomic analysis of RPAP3-GFP IPs. The subunits with a grey box are subunits not detected in the RPAP3-GFP IP.

RPAP3 is not part of the mature INO80, SRCAP and TIP60 complexes. Yet, we previously showed that RPAP3 binds co-translationally to the large subunits of these complexes: SRCAP, EP400 and BRD8^25^. We were thus interested in determining at which point RPAP3 protein dissociates from these complexes. To this aim, we performed an AP-MS analysis of GFP-RPAP3 with and without CB-6644 treatment using HeLa cells expressing a GFP-RPAP3 fusion (Figure 7C, Table S8). Remarkably, while we detected little association of RPAP3 with subunits of INO80, TIP60 and SRCAP in untreated conditions, a strong binding of RPAP3 to EP400 and ACTL6A was observed after 16h of CB-6644 treatment (2 to 4-fold increase; Figure 7C). Taken together, these results thus suggested that RPAP3 binds co-translationally to EP400 and remains associated with this subunit until RUVBL1/RUVBL2 hydrolyzes ATP. Consistently, RUVBL1/RUVBL2 are associated with ADP in the mature form of INO80, SRCAP and TIP60 complexes ^51,52,53^.

### The RUVBL1/RUVBL2 DII domain is required for their high affinity binding to RPAP3

The data above suggested that the nucleotide status of RUVBL1/RUVBL2 may directly or indirectly contribute to RPAP3 binding and dissociation, particularly during the biogenesis of INO80, TIP60 and SRCAP. To explore this possibility, we performed *in vitro* binding assays using SPR and recombinant proteins (Figure 8). We previously demonstrated that the C-terminal domain of RPAP3 interacts with the RUVBL1/RUVBL2 ring, at the top of the ATPase core and opposite to domain II (DII; ^15^). To gain a more comprehensive understanding of the molecular determinants governing RPAP3 binding, we applied the Extract2Chip (E2C) approach ^54^ to measure the *in vitro* binding affinity of native full-length RPAP3 (herein named E2C-RPAP3) to RUVBL1/RUVBL2. Indeed, in this assay, RPAP3 is immediately immobilized on the SPR chip following cell lysis. We first assessed its interaction with wild-type RUVBL1/RUVBL2 and then compared this to its binding to ATPase-deficient mutants and to RUVBL1/RUVBL2 complexes lacking the DII domain. SPR analyses revealed that E2C-RPAP3 binds RUVBL1/RUVBL2 with high affinity (Figure 8A and S5E). The isolated C-terminal domain of RPAP3 interacted with RUVBL1/RUVBL2 with a K_D_ of 4.2 nM, and E2C-RPAP3 showed a comparable affinity (K_D_ of 1.7 nM). Notably, no major differences in binding affinity were observed when SPR experiments were performed using an ATPase-deficient RUVBL1/RUVBL2 mutant purified in the presence of saturating ATP concentrations (Figure 8B and S5E). This mutant, which retains ATP binding but is unable to hydrolyze ATP and is therefore locked in an ATP-bound conformation, interacted with E2C-RPAP3 similarly to the wild-type complex. In contrast, RUVBL1/RUVBL2 complexes lacking the DII domain exhibited reduced affinity toward E2C-RPAP3, as indicated by the transient interaction profile (Figure 8C and S5E). Structural studies have shown that the RPAP3 C-terminal domain, interacting with the ATPase core, is preceded by a long unstructured region hanging along the outer face of the RUVBL1/RUVBL2 ring, connecting the TPR and C-terminal domains. In this context, our data suggest that while conformational changes of the DII domain associated with the ATPase cycle do not measurably impact E2C-RPAP3 binding affinity, the presence of the DII domain is required to further stabilize the interactions involving multiple regions of E2C-RPAP3, including those beyond the C-terminal domain. Loss of DII likely disrupts this structural framework, resulting in weakened and more transient association of E2C-RPAP3. Possibly, conformational changes of the DII during the assembly of the chromatin remodeling complexes, or its association with client subunits, may trigger RPAP3 release. However, we cannot exclude the possibility that an interaction partner (e.g. molecular chaperone) in complex with E2C-RPAP3 may modulate the binding profile with the DII deleted RUVBL1/RUVBL2 variants.

**Figure 8.**
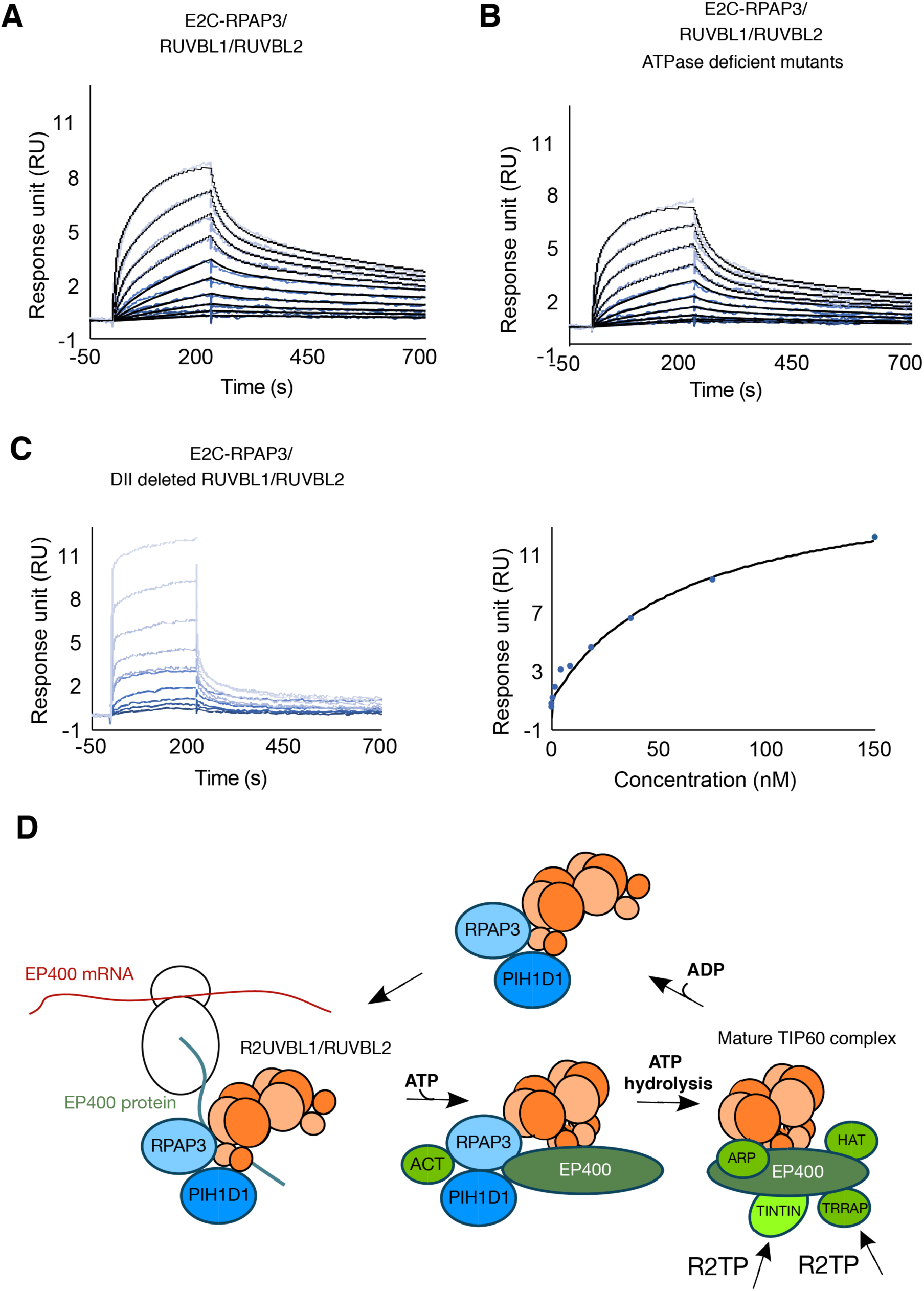
Kinetic characterization of the interaction between RPAP3 and RUVBL1/RUVBL2 complexes and model for the assembly of TIP60 complex by R2TP. **(A-C)** RPAP3 was immobilized by capturing its biotinylated Avi-tag using NeutrAvidin™-coated CM5 sensor chips. **(A)** Injection of wild-type RUVBL1/RUVBL2 at 10 different concentrations, generated by 1:2 serial dilutions with the highest concentration of 10 nM. The association rate (k_a_ = 3.5E^+7^) and dissociation rate (k_d_ = 6.2E^-2^) as well as the dissociation constant (K_D_ = 1.7E^-9^) were determined. **(B)** Injection of ATPase-deficient mutants of RUVBL1/RUVBL2 at 10 different concentrations, generated by 1:2 serial dilutions with the highest concentration of 10 nM. The association rate (k_a_ = 1.8E^+7^) and dissociation rate (k_d_ = 3.0E^-2^) as well as the dissociation constant (K_D_ = 1.7E^-9^) were determined. The kinetic profile resembles the wild-type profile **(C)** Injection of DII-deleted RUVBL1/RUVBL2 at 10 different concentrations, generated by 1:2 serial dilutions with the highest concentration of 150 nM. The dissociation constant (K_D_ = 6.1E^-8^) was determined in steady state mode (insertion to the right). The kinetic profile differs from that of wild-type and ATPase-deficient mutants RUVBL1/RUVBL2, and the calculated K_D_ suggests that the DIIs contribute to a high affinity interaction between RPAP3 and RUVLB1/RUVBL2. **(D)** Scheme illustrating the assembly of the TIP60 complex by R2TP. ARP, TINTIN, HAT et TRRAP are others sub-complexes contained in TIP60 ^51^. A similar pathway is likely used by IN080 and SRCAP.

## Discussion

In this work, we combined a series of high and medium-throughput methods to explore the cellular functions of the R2TP assembly chaperone. This validated ∼115 new RPAP3 and PIH1D1 interaction partners whose expression often depended on HSP90 and R2TP activity, and which could thus be new client proteins. Surprisingly, our results show that the INO80, SRCAP and TIP60 chromatin remodeling complexes are genuine R2TP clients, which have the unusual feature of keeping RUVBL1/RUVBL2 as resident chaperone subunits. Altogether, our results explain how cells build these essential cellular machines and clarify the role of the R2TP-specific subunits during the chaperone cycle of RUVBL1/RUVBL2, as they assemble client complexes.

### RPAP3 is required for the co-translational loading of RUVBL1/RUVBL2 on clients, including INO80, SRCAP and EP400

To clarify the role of RPAP3 in the functions of RUVBL1/RUVBL2, we used the dTAG rapid depletion system and analyzed the protein and RNA interactomes of RUVBL1 with and without RPAP3. Remarkably, SRCAP, EP400 and BRD8 were among the top 4 mRNAs whose association with RUVBL1 was most decreased in absence of RPAP3 (4-8-fold loss after a 2h depletion). Because we previously showed that the same three mRNAs are among the top ones that associate co-translationally with both RUVBL1 and RPAP3 ^25^, these results demonstrate that RPAP3 plays a direct and essential role in the co-translational loading of RUVBL1/RUVBL2 on the SRCAP, EP400 and INO80 proteins. Interestingly, these proteins are structurally related and are the motor and scaffolding subunits of their respective complexes (SRCAP, TIP60 and INO80). Moreover, RUVBL1/RUVBL2 directly and extensively interact with them, in large part via their long ATPase ‘insertion domain’ that runs through the interior of the closed DII domains of the RUVBL1/RUVBL2 hexamer, forming an unusually complex topology sometimes called the ‘spoked wheel’ ^51,52,55,56^ (for a reviemw, see^37^). Given that the DII domains of RUVBL1/RUVBL2 can make large scale movements and switch between open and closed conformations ^20,22^, the function of RPAP3 may be to keep RUVBL1/RUVBL2 in an open conformation compatible with binding, before the DII domains close on the ‘insertion domain’ and lock the complex (Figure 8D, see below). Additionally, RPAP3 strongly associates with INO80 in LUMIER co-IP experiments. It is thus also possible that RPAP3 also directly binds INO80, SRCAP and EP400 to facilitate the loading of RUVBL1/RUVBL2.

It is interesting to note that the binding of RUVBL1 to INO80 mRNA is ∼10 fold less than SRCAP and EP400 mRNAs ^25^. Like SRCAP and EP400, INO80 interacts with RUVBL1/RUVBL2 via a long ‘insertion domain’ that runs through the hexameric DII domains ^51,56,57^. However, the ‘insertion domain’ is located much closer to the C-terminal end for INO80 than SRCAP or EP400 (300 amino-acids vs 1200, ^51,56,57^). This is reminiscent of the situation observed for PIKKs, where RUVBL1 associates co-translationally only with SMG1 ^25^, which contains an additional 1000 amino-acid C-terminal domain, as compared to the other PIKKs ^25,58^. This suggests that the ribosome dwell time and the location of protein interaction domains are key determinants of RUVBL1/RUVBL2 co-translational binding. In the future, it will be interesting to investigate whether ribosome pausing facilitates co-translational interactions, or conversely, whether the lack of co-translational interaction induces pausing, eventually triggering degradative quality control pathways and enabling the co-regulation of protein partners.

The role of RPAP3 in promoting the co-translational binding of RUVBL1/RUVBL2 extends to other R2TP clients and appears to be a general function of this protein, as RPAP3 depletion led to the loss of RUVBL1 co-translational binding to the large RNA polymerase subunits, AGO proteins, PCF11/SCAF family and CHD chromatin remodelers (Table S7). Interestingly, CHD proteins share a similar fold as INO80, SRCAP and EP400, but generally function in simpler complexes or as isolated proteins, and never have RUVBL1/RUVBL2 as resident proteins ^59,37^. Possibly, RPAP3 and RUVBL1/RUVBL2 played an ancient and transient role in the biogenesis of ATPase chromatin remodelers, and during evolution became permanently captured in the INO80/SRCAP/EP400 family, thanks to their ‘insertion domain’ and their increased in number of subunits.

### RPAP3 is required for the integrity of INO80, SRCAP and TIP60 complexes

The role of RPAP3 in the biogenesis of INO80, SRCAP and TIP60 likely extends beyond the loading of RUVBL1/RUVBL2 on the motor subunits. Indeed, mass-spectrometry analyses of RUVBL1 and INO80 IPs show a severe disruption of the INO80, SRCAP and TIP60 complexes after the depletion of RPAP3, with the loss of many, but not all, subunits of these complexes. Interestingly, RUVBL1 remains associated with the EP400, SRCAP and INO80 motor subunits, which in contrast show the highest loss at the co-translational level. This is likely because co-translational interactions at the mRNA levels reflect early events in the biogenesis of these proteins, while the proteomic analysis reflects later assembly steps and also includes some of the pre-existing complexes in which RUVBL1/RUVBL2 are already stably associated with the EP400, SRCAP and INO80 proteins. Yet, the fact that multiple subunits are lost upon RPAP3 depletion suggests that it plays additional roles beyond loading RUVBL1/RUVBL2 (see model in Figure 8D). The analysis of subunit loss with respect to the structure of these complexes shows that RPAP3 depletion leads to the loss of the TINTIN module of TIP60, which is composed of MORF4L1, MORF4L2, MRGBP and BRD8 ^51^. Our Y2H and LUMIER co-IP assays show that RPAP3 is likely a direct partner of MORF4L2 and additionally, RPAP3 and RUVBL1 co-translationally associate with BRD8 ^25^. This suggests that the R2TP may directly associate with the TINTIN module of TIP60 to promote its assembly with EP400, possibly independently of the RUVBL1/RUVBL2 associated with EP400 itself, as these do not make strong contacts with the TINTIN module ^51^. Another interesting case is the one of TRRAP, the PIKK-like subunit of TIP60 ^60^. Indeed, the 6 PIKKs have been shown to require R2TP for their biogenesis and incorporation in functional complexes, via an adaptor called the TTT complex ^32^. It is thus possible that the binding of TRRAP to EP400 is promoted by R2TP, again independently of the RUVBL1/RUVBL2 hexamer loaded on EP400 (Figure 8D).

### The ATPase cycle of RUVBL1/RUVBL2 controls the biogenesis of INO80, SRCAP and TIP60 complexes

The RUVBL1/RUVBL2 ATPases in the mature INO80 and SRCAP complex are in an ADP form^39,52,56^, and this is also likely the case for TIP60 ^61^. In addition, we previously showed that the inhibition of the ATPase activity of RUVBL1/RUVBL2 with the CB-6644 inhibitor, which blocks them in an ATP form, prevents their co-translational binding to SRCAP, EP400 and INO80 mRNAs ^25^. This suggests a step-wise assembly pathway regulated by the nucleotide status of RUVBL1/RUVBL2 (Figure 8D). First, the initial co-translational binding of RUVBL1/RUVBL2 to their clients would occur in a form devoid of nucleotides, thus with the DII domains in an open conformation ^22^. This takes place with the RUVBL1/RUVBL2 in the full R2TP complex, as RPAP3 co-translationally binds the same mRNAs as the RUVBLs and is required for their co-translational binding. Client binding would trigger ATP loading and the closure of the DII domain on the SRCAP, INO80 and EP400 insertion domain, with RPAP3 still present at this stage as evidenced by the strong binding of RPAP3 to EP400 in presence of CB-6644. The closure of the DII domains would then facilitate the binding of additional subunits and ATP hydrolysis, leading to the formation of the final mature complexes with the RUVBLs in an ADP bound, closed form. RPAP3 would be evicted from the complex during the final assembly stages, most likely because engagement of the DII domains with client/additional subunits, or conformation changes destabilizing RPAP3 binding.Eventually, RUVBL1/RUVBL2 may be recycled into R2TP by reassociating with the RPAP3/PIH1D1 heterodimer and releasing ADP. This would be consistent with recent biochemical findings indicating that the RPAP3/PIH1D1 heterodimer stimulates RUVBL1/RUVBL2 ATPase activity by enhancing nucleotide dissociation ^62^, and with structural data indicating that RPAP3/PIH1D1 modifies the conformation of nucleotide-bound RUVBL1/RUVBL2, thereby opening a pocket enabling nucleotide exchange ^20^.

INO80, SRCAP and TIP60 share the unusual property of containing RUVBL1/RUVBL2 as resident subunits. Nevertheless, the model outlined here resembles the assembly pathway of other R2TP client complexes. For instance, it has been proposed that NOP58 is loaded on R2TP in absence of nucleotide, and that ATP binding leads to the formation of a snoRNP assembly intermediate devoid of RPAP3 and PIH1D1, but containing the snoRNA, SNU13 and the assembly factors ZNHIT6, NUFIP1 and ZNHIT3^33^. The resolution of this assembly intermediate requires ATP hydrolysis ^33^, which leads to the release of the snoRNP assembly factors with RUVBL1/RUVBL2 in an ADP form. Thus, a key difference with the INO80 type of complexes is that upon ATP hydrolysis, RUVBL1/RUVBL2 remain on the client in the case of INO80, while they remain with the assembly factors for C/D snoRNPs. In the future, it will be interesting to determine whether assembly factors for C/D snoRNP or other R2TP clients possess insertion-like domains mediating association with RUVBL1/RUVBL2. More generally, the rich structural data available on INO80, SRCAP and TIP60 may help understand how RUVBL1/RUVBL2 scaffold the assembly of client complexes, possibly providing a unifying mechanism for R2TP clients. The large interaction resources of this work also provide opportunities to assess new functions for the R2TP chaperone, in particular as many of the partners found are in complexes not previously known to be assembled by R2TP.

## Material and methods

### Cell culture and generation of CRISPR/Cas9 clones

HEK293 and HEK293T cells were grown in DMEM containing 10% fetal bovine serum and 1% penicillin/streptomycin, at 37°C, 5% CO_2_. HCT116 cells were grown in McCoy’s 5A instead of DMEM. HCT116 cell line have been previously edited by CRISPR/Cas9 to contain homozygous insertion of a mini-AID-3xHA-IRES-Hygromycin cassette in RPAP3 gene and a TIR1 expression gene ^31^. This cell line and parental cells were further knock-in to insert a GFP-IRES-Neomycin cassette at the C-terminus of the endogenous INO80 gene. We also used HCT116 cell line with a GFP-IRES-Neomycin cassette inserted in fusion with the C-terminus of the RUVBL1 gene ^25^ which was further edited with a 3HA-dTAG-IRES-Puromycin cassette inserted homozygously at the C-termini of the RPAP3 gene. The sequences targeted by the guide RNAs were (PAM sequences are lowercase): TCAGCAGTAAGAGACTCCCCagg for *RUVBL1* guide 1, ACTTCATGTACTTATCCTGCtgg for *RUVBL1* guide 2; GAGCTGAAAAAGAGGTATGGtgg for RPAP3 guide. Clones were characterized by genotyping, Western blotting and immunofluorescence. PCR was conducted using a Phusion polymerase (NewEngland Biolabs) on genomic DNA prepared with Wizard® Genomic DNA Purification Kit (Promega). Stable HeLa cell line expressing GFP-RPAP3 were generated by Flp-In recombination and selected on Hygromycin B.

### Plasmids

For the LUMIER co-IP assays, Gateway recombination cloning was done with pcDNA5-FRT-3xFLAG-FL-Rf for the bait, and L30-HA-RL-Rf for the prey. Orfs were initially in a donor vector from the Orfeome 9 ^44^ or the Life technology orfeome library. Maps are available upon request.

### Yeast two-hybrid screen

Yeast two-hybrid screening was performed by Hybrigenics Services, S.A.S., Paris, France (http://www.hybrigenics-services.com). The coding sequence for full-length RPAP3 (NCBI reference <u>NM_024604.3</u>) was PCR-amplified and cloned into pB27 plasmid, which derives from pBTM116 as previously described ^15^. Briefly, random-primed cDNA library was constructed in derivative of the P6 plasmid pGADGH with cDNAs made from three human lung cancer cell lines: A549, H1703, and H460. As many as 51 million clones were screened. This represents 5-fold the complexity of the library. A mating strategy was used for the screens, using on one side L40ΔGal4 (Mata) yeast strains transformed with the bait, and on the other side the strain YHGX13 (Y187 ade2-101::loxP-kanMX-loxP, Matα) containing the library. His+colonies were selected on a medium lacking tryptophan, leucine and histidine. The 5′ and 3′ junctions of the prey insert were analyzed by capillary sequencing after yeast lysis and PCR amplification. The resulting sequences were compared to Human GenBank (NCBI) for prey identification. A statistical analysis of the results has been conducted to define a confidence score which identify False positive (Score F), and highly connected preys (score E). The other scores defined a probability for an interaction to be identified by chance, and are divided in four categories, from A (highest confidence, Y2H score of 8) to D (lowest confidence, given a Y2H score of 2) and B and C in between (Y2H score of 6 and 4 respectively).

### AF3 interaction screen

The prediction of the three-dimensional structures of the studied biomolecules was performed using AlphaFold 3. The sequences of interest were submitted via the AlphaFold 3 platform using default parameters. The generated models were evaluated using the confidence scores provided by the software and then visualized for structural analysis. An AF3 score was attributed according to the minimal predicted aligned error (PAE; in angstrom) over the interacting region predicted from the Y2H data, as follow: PAE 0-5 = 8; PAE 5-10 = 6 ; PAE 10-15 = 4 ; PAE 15-20 = 2; PAE 20-25 = 1.

### LUMIER co-IPs

HEK 293T cells were grown on 24-well plates and co-transfected with 450 ng of the RL fusion and 50 ng of the 3xFLAG-FL fusion and 2 μl of JetPrime (Ozyme). After 48h, cells were lysed in 450 µl of HNTG lysis buffer containing protease inhibitor cocktail (Complete, Merck), incubated for 15 min at 4°C and spun down at 4°C and at 20,000 × g for 15 min. 100 µl of the extract were dispatched in two wells of a 96-well plate (Lumitrac, Greiner), with one well coated with anti-FLAG antibody (10 µg/ml in PBS; F1804 Sigma-Aldrich), and one control well without antibodies. Both wells are blocked with 300 μl of blocking solution for 1h (5% BSA, 3% sucrose and 0.5% Tween-20 in PBS). Plates were incubated with the extract for 3h at 4°C, and then washed 5 times with 300 µl of ice-cold HNTG, for 10 min at 4°C for each wash. After the last wash, 10 µl of PLB (Protein Lysis Buffer, Promega) was added in each well. To measure the signal in the input, 2 µl of extract and 8 µl of PLB was put in empty remaining wells. Plates were then incubated 5 min at room temperature, and FL and RL luciferase activities were measured in IP and input wells, using the dual luciferase kit (Promega). For each prey, co-IP efficiency was defined as the RL IP/Input divided by the FL IP/Input. When three replicates were available, statistical significance was evaluated using t-test assaying whether the co-IP efficiency in the anti-FLAG IP was greater than the values obtained in the control IP, done without antibodies. When the pvalue was lower than 0.05 and the enrichment fold over the control IP was larger than 5, the prey was considered as true interactant. When the assay was done in a single replicate, only preys giving values with a enrichment fold greater than 15 were considered as true interactants.

### Direct interaction score calculation

First, for each prey, we attributed a LUMscore depending on the LUMIER co-IP efficiency: >2% co-IP efficiency yielded a score of 8; 0.8-2%: 6; 0.32-0.8%: 4; 0.128-0.32%: 2; <0.128%: 1. Proteins not validated in LUMIER IP (false interactants, see above) had a score of 0, irrespective of their co-IP efficiency. Then, we calculated a DI Score (Direct Interaction Score) by adding the LUMscore, Y2Hscore and AF3score (respectively obtained from Y2H screen and AF3 analysis respectively, see above).

### Functional luciferase assay

HEK 293T cells were grown on 24-well plates and co-transfected with 450 ng of the RL fusion and 50 ng of the 3xFLAG-FL and 2 μl of JetPrime (Ozyme). After 48 h, cells were lysed in 150 μl of passive lysis buffer (Dual luciferase assay kit, Promega) and incubated at 4°C for 15 min. RL and FL activities were measured on 96-well plates using 8 μl of cell extract with 50 μl of each solution of the dual-luciferase assay kit (Promega). Values obtained for RL were normalized to FL values obtained in the same well. Experiments were done in triplicate. For treatment with geldanamycin, drug was added for 16h at 2 μM before cell lysis. For depletion of PIH1D1, experiments were done in HEK 293T CRISPR-KO for PHID1 ^14^ in parallel to parental cells. For depletion of RPAP3, experiment was done in HCT116 having a mini-AID-3xHA-IRES-Hygromycin cassette in RPAP3 gene parallel with HCT116 and a TIR1 expression gene ^31^ in parallel with HCT116 cells. A luciferase assay with RL fusion to mPHAX was also done in all the conditions to have normalization between the different cellular conditions used.

### Genetic interaction mapping screen

Genetic interactions screens for PIH1 deletion strains were performed as previously described^46,65^. Briefly, the query strains were generated from deletion strains bearing the KanMX4 marker by replacement with a haploid specific nurseothricin resistance cassette. Mass mating with a pooled library of deletion and DAmP^66^ (decreased abundance by mRNA perturbation) strains was followed by sporulation and selection for double mutant spores. Growth of individual strains was estimated by custom Agilent microarray hybridization for the specific barcodes that mark each mutant in comparison with a control experiment. The obtained query/reference fitness values were normalized and corrected for pleiotropy effects^46^. Normalized results were expressed as log2(Q/R) where Q represents the signal intensities of a given mutant when combined with the query allele (*pih1*Δ), and R is the signal from the same mutant when introduced in a reference population. Negative and positive log2(Q/R) values represent aggravating and buffering (or alleviating) effects, respectively^47^.

### Proteomic analysis of SILAC IP and label-free AP-MS

For SILAC IP experiments, HCT116 INO80-GFP cells with or without mAID-RPAP3 were grown for 15 days in each isotopically labeled media (CIL/Eurisotop), to ensure complete incorporation of isotopically labeled arginine and lysine as follow: light label (R0K0, L); semi-heavy label with L-Lysine-2HCl (2H4) and L-Arginine-HCl (13C6) (R6K4, M); heavy with L-lysine–2HCl (13 C6; 15 N2) and L-arginine–HCl (13 C6; 15 N4) (R10K8, H). Six 15-cm diameter plates were used per SILAC condition. Cells were treated with 500 μM of a derivative of auxin (Indole-3-acetic acid sodium salt, I5148 Sigma Aldrich) for 16h before lysis. Cells were rinsed with PBS, trypsinized and cryogrinded^67^ and the powder was resuspended in HNT lysis buffer (20 mM HEPES, pH 7.4, 150 mM NaCl, 0.5% triton X-100, protease inhibitor cocktail (Roche). Extracts were incubated 20 min at 4°C and clarified by centrifugation for 10 min at 20,000 x g. Extracts were pre-cleared by incubation with Protein G Sepharose beads (GE healthcare) for 1h at 4°C. The control was extracted from the SILAC light condition prepared from parental HCT116 cells that did not express the GFP fusion. Each extract was then incubated with 50 µl of GFP-Trap agarose beads (Chromotek) for 1.5h at 4°C, washed 5 times with HNT buffer, and beads from the different isotopic conditions were finally pooled. Bound proteins were eluted by adding 1% SDS to the beads and boiling for 10 min. Proteomic analysis was performed as previously described ^15^.

For label-free proteomic analysis, HCT116 RUVBL1-GFP cells having a dTAG on the two RPAP3 alleles were seeded in 15 cm plate. 2h before lysis, dTAG-V1 was added at 50 nM final. Cells were lysed with 0.5 ml HNTG lysis buffer (20 mM HEPES, pH 7.9, 150 mM NaCl, 1% Triton X-100, 10% glycerol, 1 mM MgCl2, 1 mM EGTA) supplemented with proteinase inhibitor cocktail (Roche). Cells were scraped and incubated for 30 min at 4°C before being centrifuged for 10 min at 16000g at 4°C. The supernatant was subjected to immunoprecipitation using GFP-Trap agarose beads (Chromotek). Agarose control beads were used for non-specific control IP. Extracts were incubated with beads for 1.5h at 4°C and washed 5 times with ice-cold HNTG. Proteins were then eluted in Laemmli buffer and heated 10 min at 95°C. Sample treatments for mass spectrometry analysis were already described ^25^. All experiments were done in triplicates.

#### RIP-Seq analysis

HCT-116 RUVBL1-GFP cells with or without RPAP3-dTAG were seeded in a 15 cm plate. 2h before lysis, dTAG-V1 was added at 50 nM final. At 70% confluency, cells were put on ice and washed twice with ice-cold PBS, and then lysed with 0.5 ml HNTG lysis buffer supplemented with a cocktail of protease inhibitors (Roche). Cells were incubated for 30 min on a tube roller at 4°C before being centrifuged for 10 min at 16000g at 4°C. The supernatant was subjected to immunoprecipitation for 1.5h at 4°C using GFP-Trap agarose beads (Chromotek). Agarose control beads were used for non-specific control IP. All experiments were done in duplicates. Beads were washed 5 times with ice-cold HNTG. RNeasy mini kit (Qiagen-74106) was used to purify RNAs. DNA contamination was eliminated by a TURBO-DNA Free kit, as recommended by the manufacturer (Invitrogen-AM1907).

SMART-Seq Stranded Kit (Takara Bio) was used to generate cDNA libraries for high throughput sequencing using NovaSeq 6000 Illumina. RNA and DNA libraries quality were subjected to quality control checks periodically (or after each step) using a fragment analyzer (Kit high sensivity NGS) and qPCR (Roche Light Cycler 480). Ribosomal cDNAs were cleaved with scZapR enzyme in the presence of scR-Probes designed against mammalian ribosomal RNAs. Libraries were then sequenced with NovaSeq Reagent Kits on a flow cell SP (single read -100 nucleotides) using the 2-channel sequence by synthesis method. Before demultiplexing, sequences were mixed with PhiX sequences as an Illumina control (not indexed). Demultiplexing was realized using Illumina bcl2fastq software (v2.20.0.422) and all PhiX sequences were removed.

FastaQC software (v0.11.9) provided a tool to check read quality after high-throughput sequencing and MultiQC program (V1.12) served to group data that emerges from all samples. FastQ screen software served to detect contaminating genomes. Sequences were aligned on 15 species genomes, mycoplasma, and ribosomal RNAs using Bowtie2 software. No contamination was detected.

TopHat2 software (v2.1.1) was used to align the obtained reads on the human genome (hg38 version) and to annotate them according to NBCI database. FeatureCounts software was utilized to count reads per gene. All reads that overlap on two genes or show alignment at different positions on the genome were discarded. Counts were normalized in each sample to eliminate differences due to manipulation. This was done by a normalization method known as relative log expression (RLE). Differential expression genes were identified using EdgeR (v3.34.1). Tests are based on a generalized linear model and allow counts modeling using a negative binomial distribution. To calculate the reads per kilobase per million mapped reads (RPKM), the formula below was used: (read counts aligned on a specific gene/ (total aligned counts x gene length) x109.

### Production of RUVBL1/RUVBL2 complexes, and cellular biotinylated RPAP3

The coding sequences for His_6_-Thrombin-RUVBL1(1-456) and RUVBL2(1-463)-HRV3C-FLAG-Fh8 were synthesized and cloned in pETDuet^TM^-1 in MCS 1 and 2, respectively, by GenScript. ATP hydrolysis-deficient (His_6_-Thrombin-RUVBL1(1-456)_D302N and RUVBL2(1-463)_D299N-HRV3C-FLAG-Fh8) and DII-truncated (His_6_-Thrombin-RUVBL1(1-126; 234-456) and RUVBL2(1-133; 238-463)-HRV3C-FLAG-Fh8) variants of the RUVBL complex were synthesized and cloned in the same manner as the wild-type construct. The coding sequences for BirA-FLAG and AviTag-His_6_-RPAP3(1-665) were synthesized and cloned into pRSF-Duet^TM^-1 in Multiple Cloning Site (MCS) 1 and 2, respectively, by GenScript. The coding sequence for AviTag-His_6_-PIH1D1(1-290) was synthesized and cloned into pRSF-Duet^TM^-1 in MCS 2, replacing the AviTag-His_6_-RPAP3(1-665) coding sequence, by GenScript. The coding sequences for BirA-FLAG, AviTag-His_6_-RPAP3(1-665) and His_6_-PIH1D1(1-290) were synthetized and cloned into a customized version of pET-Duet^TM^-1, referred as pET-Triet, by Genscript. This modified vector was engineered by introducing a third MCS, based on MCS1, downstream of the terminator sequence of MCS2. Production of RUVBL1/RUVBL2 complex wild-type, ATP hydrolysis-deficient and DII-truncated variants were previously performed ^68^.

For cellular biotinylation of the RPAP3 protein, the DNA plasmids containing BirA-His_6_ and AviTag-His_6_-RPAP3 coding sequences were transformed in BL21 Star™ (DE3) pRARE2 and grown overnight at 37°C in Luria-Bertani (LB) agar plates with the respective antibiotics as selection agents. Fresh colonies were picked and grown overnight (ON) at 37°C and 150 rpm in Power Broth (PB) media supplemented with the respective antibiotics. Each ON culture was diluted in PB media supplemented with the respective antibiotics to a final optical density (OD) of 0.1 and grown at 37°C and 150 rpm until reaching an OD between 1.8. The proteins were expressed using 500 μM IPTG, ON at 18°C. In order to ensure cellular biotinylation of AviTag-His6-RPAP3 protein, 5 μg/mL D-Biotin was added to the cells upon IPTG addition. Finally, the cells were harvested by centrifugation at 7030 *g* and 4°C, for 15 min, and stored at - 80°C.

The cells were resuspended in BugBuster^®^ Protein Extraction Reagent (Novagen) supplemented with 0.1 mg/mL Lysozyme, 5 U/mL Benzonase, 1 mM PMSF and PIC without EDTA and placed on ice for 20 min. The lysate was cleared by centrifugation at 13200*g* and 4°C for 25 min and injected onto a PD-10 Desalting column equilibrated in buffer A (20 mM HEPES pH 8.0, 150 mM NaCl, 2 mM MgCl_2_, 0.5 mM TCEP, 1 mM PMSF and PIC without EDTA) to remove the excess of D-Biotin.

### Surface plasmon resonance (SPR) experiments

The interactions between the three versions of RUVBL1/RUVBL2 complex and the cellular biotinylated RPAP3 protein were assessed by SPR at 25 °C using a Biacore T200 instrument (Cytiva). The surface of a CM5 sensor chip (Series S) was activated with 400 mM 1-ethyl-3-(3-dimethylaminopropyl)-carbodiimide (EDC) and 100 mM N-hydroxysuccinimide (NHS) for 10 min and coated with NeutrAvidin™ Biotin-Binding Protein (Thermo Scientific, 31000) at 50 μg/mL in 10 mM sodium citrate pH 4.5. The cellular biotinylated RPAP3 at 3 mg/mL in HBS-P+ (10 mM HEPES pH 7.4, 150 mM NaCl and 0.05 % Tween® 20) was coupled to the NeutrAvidin™ Biotin-Binding Protein coated surface for 180 s at a flow rate of 10 μL/min to reach 80-85 RU. The background buffer used during immobilization was HBS-P+. The RUVBL1/RUVBL2 complex wild-type, ATP hydrolysis-deficient and DII-truncated variants were then injected over the captured RPAP3 surface at 10 different concentrations using a 2-fold dilution series, with the highest concentration of 10 nM (for wild-type and ATP hydrolysis-deficient) and 150 nM (for DII-truncated). The running buffer used during the experiment contained 20 mM NaKPi pH 7.5, 150 mM NaCl, 5 mM MgCl_2_, 1 mM DTT and 0.05 % Tween-20. Interaction analysis cycles consisted of a 220 sec sample injection followed by 650 sec of running buffer.

All sensorgrams were processed by first subtracting the binding response recorded from the control surface (reference spot), followed by subtraction of the buffer blank injection from the reaction spot. The datasets were fitted to a Heterogeneous Ligand model to determine the kinetic rate constants using the provided Biacore T200 evaluation software. The interaction affinity was additionally calculated at steady state (K_Dss_) when applicable using the provided Biacore T200 evaluation software.

## Acknowlegdments

We acknowledge (i) the sequencing facility MGX, which obtained financial support from France Génomique National infrastructure, funded as part of the “Investissement d’Avenir” program (contract ANR-10-INBS-09); (ii) the imaging facility MRI, member of the national infrastructure France-BioImaging (https://ror.org/01y7vt929) supported by the French National Research Agency (ANR-24-INBS-0005 FBI BIOGEN); and (iii) the Montpellier Proteomics Platform (PPM, BioCampus Montpellier), member of the national Proteomics French Infrastructure (ProFI UAR 2048) supported by the French National Research Agency (ANR-24-INBS-0015). The work was supported by grants from the ANR (ANR-23-CE12-0022), the INCa (PLBIO23) and the ‘Ligue Nationale contre le Cancer’ (équipe labélisée LIGUE 2023). YA and MP were supported by INCa. We thank Sylvain Tartier, Claire Abéza and Valentine Méric for their technical help in the experiments.

## Author contributions

CV and EB conceived and supervised the work. YA prepared the collection of vectors with all the orfs and performed the functional and LUMIER screens in human cells with the help of CV. YA, MP and MCR prepared the cell lines by CRISPR/Cas9. MP and SB performed the label-free proteomic and RIP-Seq experiments. CV with FV performed the SILAC proteomic experiments. EB conducted the AF3 analysis. LD and CS performed the genetic screen in yeast. AP, PB, PS and TB did the SPR experiments. SU and MS did the mass spectrometry analysis and processed the data. CB and JI performed the RNA sequencing and analyzed the data. CV and EB wrote the manuscript and prepared the figures.

## Supplementary figures and legends

**Figure S1.**
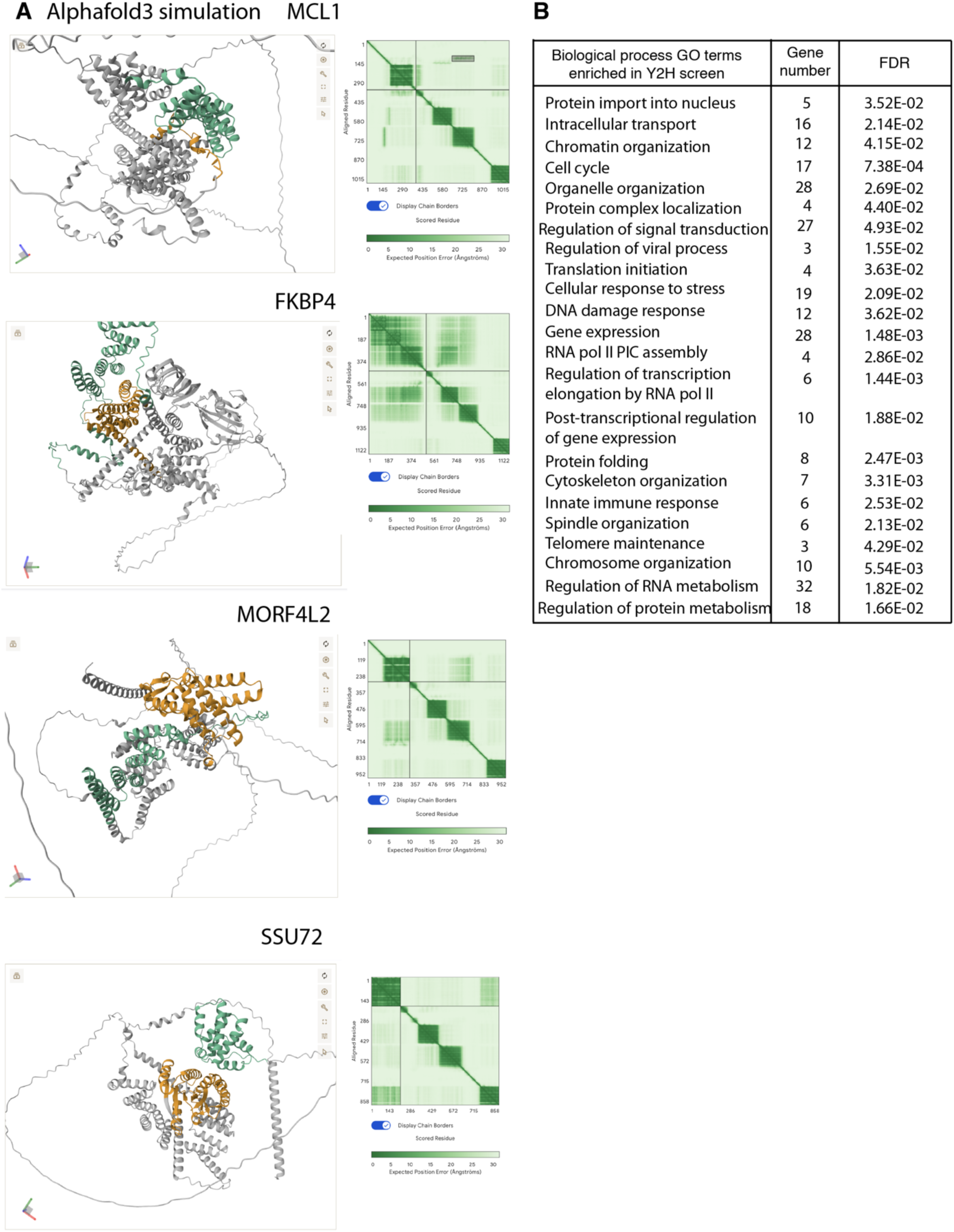
Analysis of the preys found in the RPAP3 Y2H screen. **(A)** Alphafold3 (AF3) models of the interaction of RPAP3 with MCL1, FKBP4, MORF4L2 and SSU72. Predicted interaction surfaces between candidate protein and RPAP3 are highlighted in green and orange, respectively. The left table shows the distance error between pairs of amino-acids, as estimated by AF3. **(B)** GO term enrichment analysis for Biological Processes, for the 98 preys found in the yeast two-hybrid screen. The number of genes counted in each category and their false discovery rate (FDR) are indicated.

**Figure S2.**
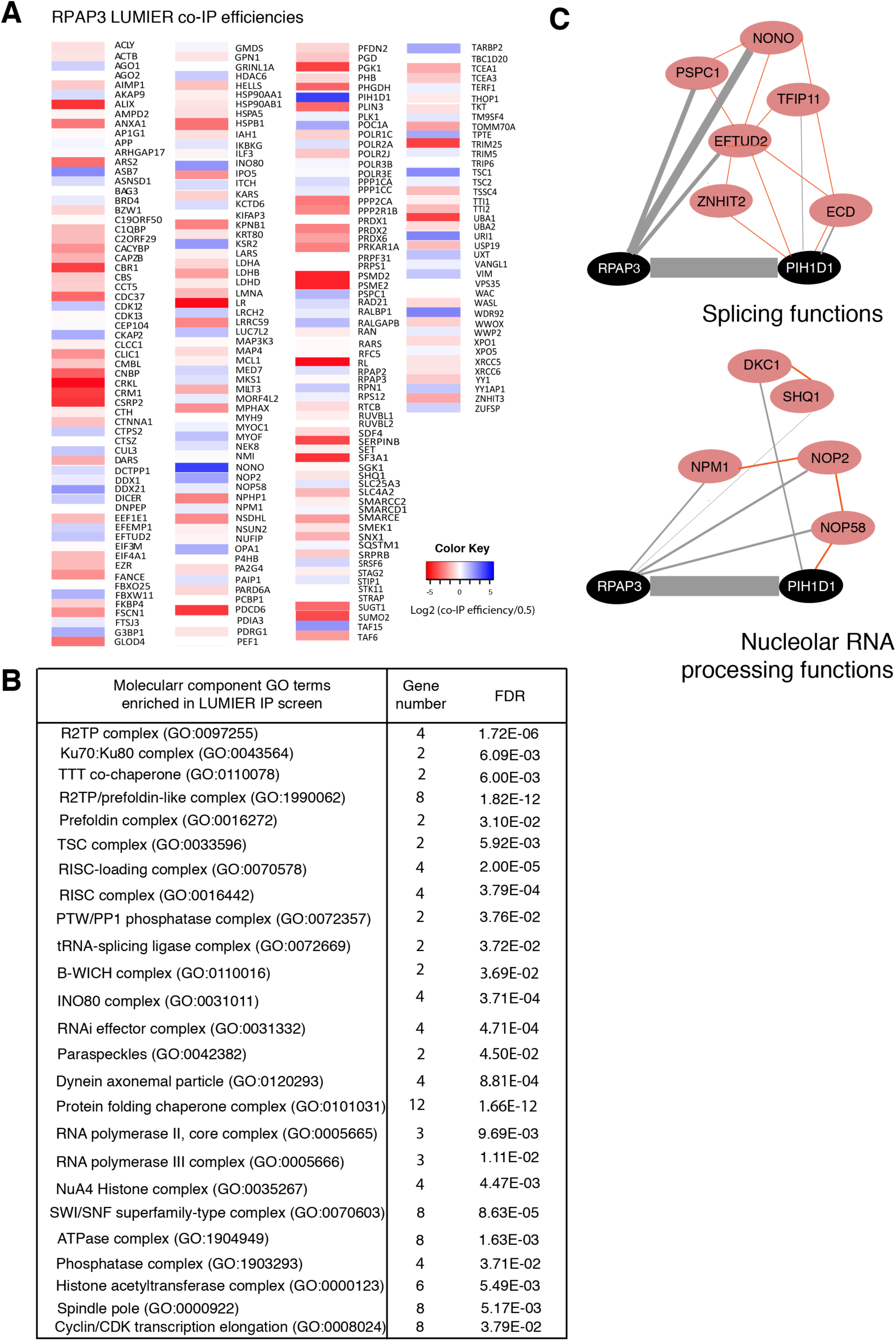
Analysis of RPAP3 and PIH1D1 interaction partners. **(A)** Heat map of LUMIER co-IP efficiency for the 228 candidate RPAP3 interaction partners. The bait was 3xFLAG-FL-RPAP3 and Alix is a control prey protein. LUMIER co-IP efficiencies were averaged from three experiments and the color represents the Log_2_(co-IP efficiency/0.5), according to the color scale shown. **(B)** GO term enrichment analysis for Molecular Components, for the RPAP3 and PIH1D1 interactants validated by LUMIER co-IP (74 and 26 interactants, respectively). The number of genes counted in each category and their false discovery rate (FDR) are indicated. **(C)** Selected network of proteins interacting with RPAP3 and PIH1D1, and involved in splicing (top) and nucleolar RNA processing (bottom) functions. Red line: interactions documented in String and in the literature.

**Figure S3.**
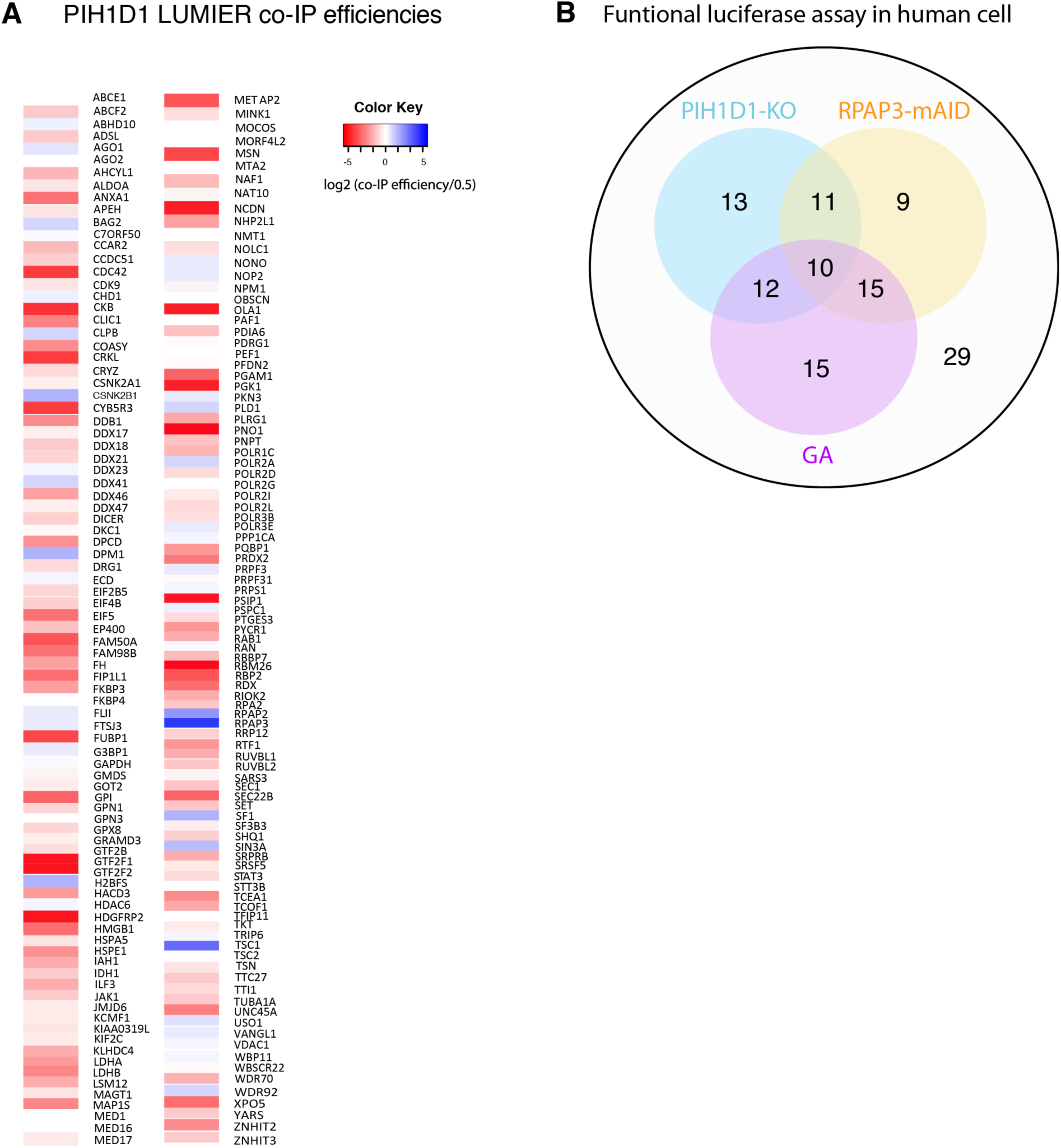
LUMIER co-IP analysis of candidate PIH1D1 interactants and effect of R2TP inhibition on client expression levels in human cells. **(A)** Heat map of LUMIER co-IP efficiency for the 186 candidate PIH1D1 interaction partners. The bait was 3xFLAG-FL-PIH1D1 and LUMIER co-IP efficiencies were measured in a single replicate and the color represents the Log_2_(co-IP efficiency/0.5), according to the color scale shown. **(B)** Venn diagram showing the number of transiently transfected proteins with level changed in the treated or depleted condition compared to the control condition (geldanamycin treated in violet, PIH1D1-KO in blue and RPAP3-mAID in yellow; Table S3). Among the 114 proteins tested, 29 were not affected in any of the conditions.

**Figure S4.**
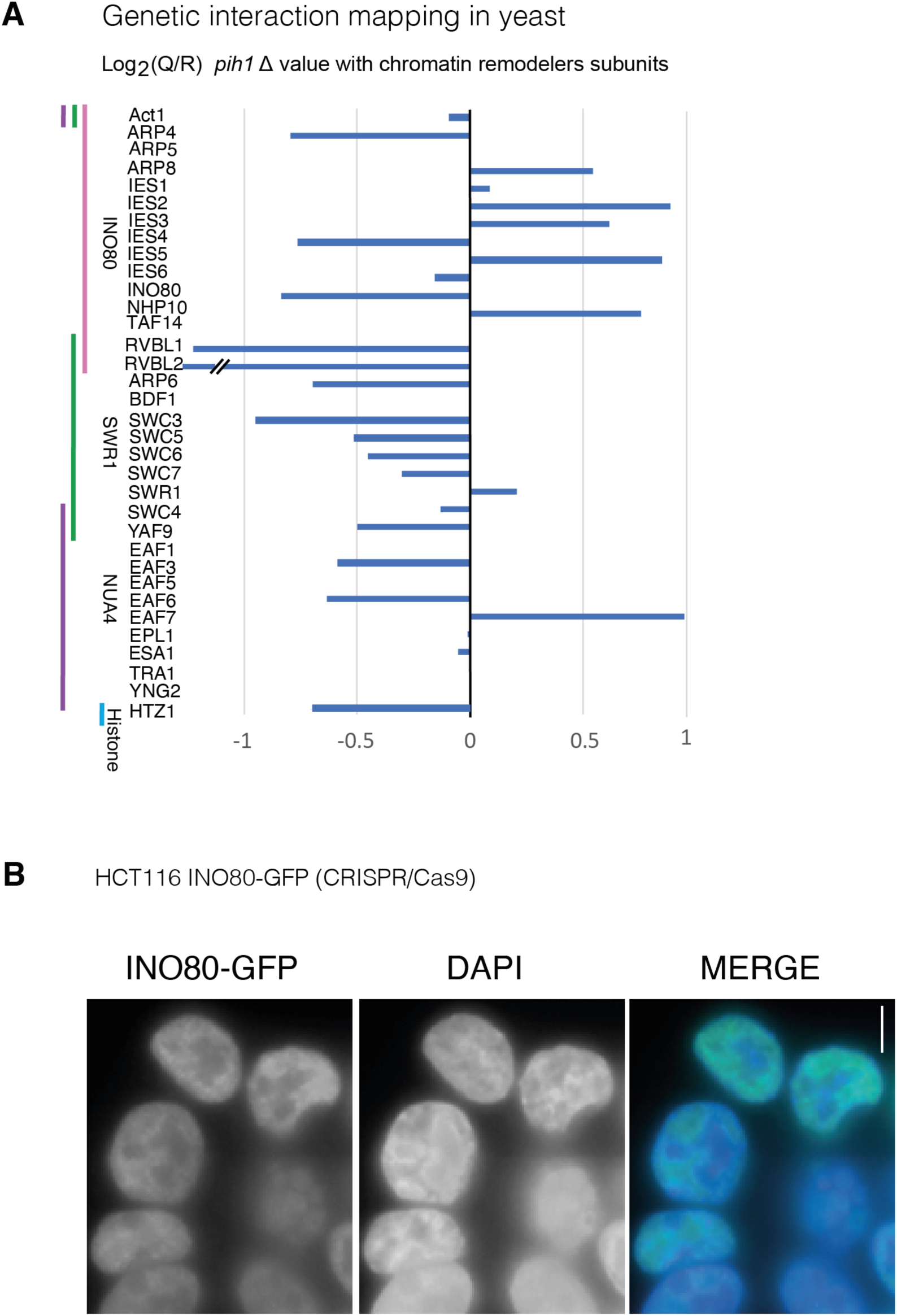
Data from genetic interaction mapping in yeast Δpih1 focused on chromatin remodeler subunits. Localization of INO80-GFP in HCT116 stable clone. (A) Barplot representing log_2_(Q/R) values obtained from the genetic screen using the deletion of PIH1 in *S. cerevisiae* only for gene names related to chromatin remodeling complex. They are ordered according to their related complex (subunits from the INO80 complex are in front of the pink line, subunits from the SWR1 complex in front of the green line and subunits from the NuA4 complex in front of the purple line). Histone HTZ1 is also represented. (B) Micrographs showing HCT116 cell edited to express INO80-GFP fusion. The GFP signal is shown on the left panel. DNA is stained with DAPI (middle panel). The merge image shows nuclear localization of INO80-GFP (right panel). Scale bar is 10 μm.

**Figure S5.**
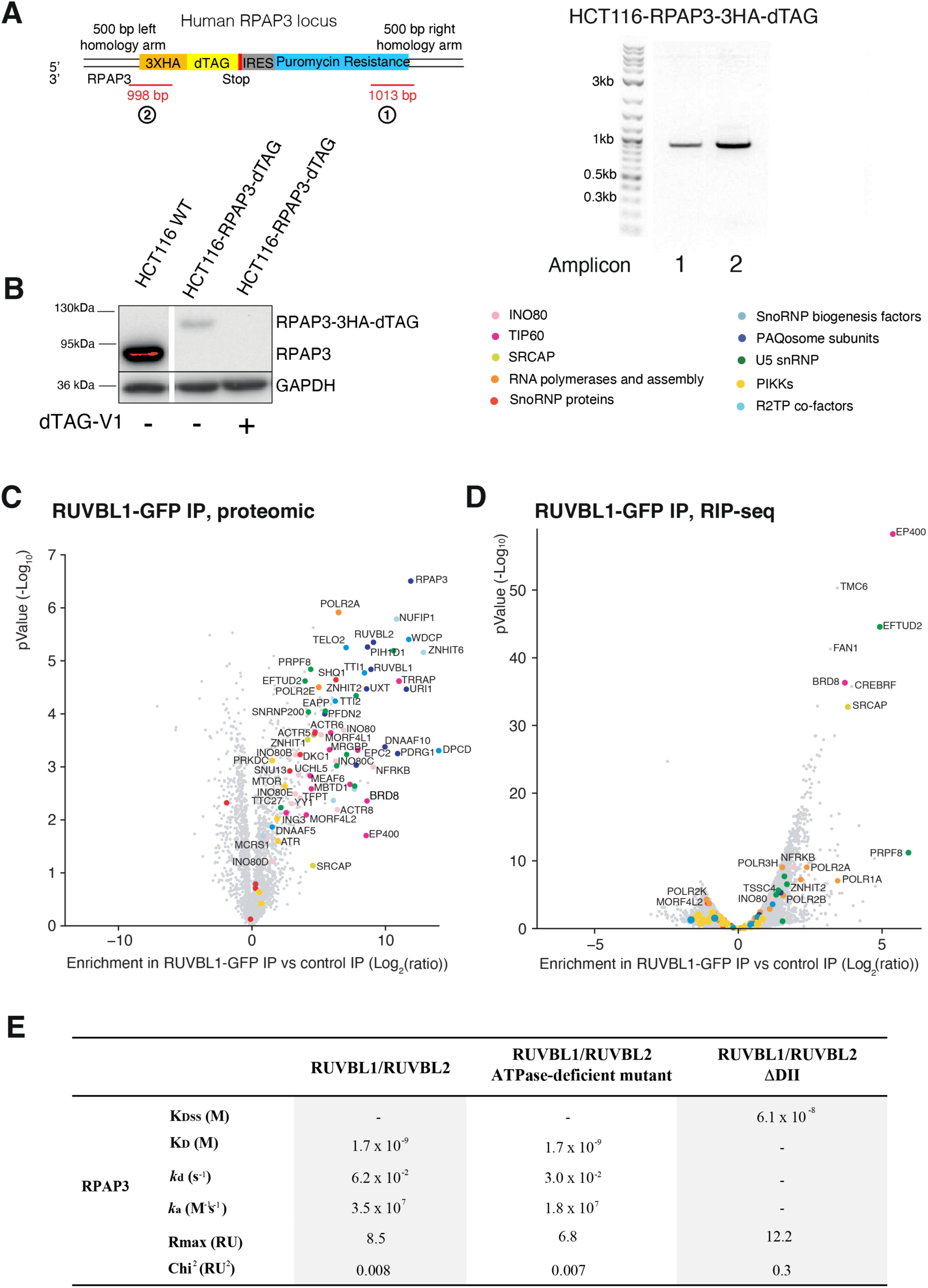
Characterization of HCT116 cells expressing RPAP3-dTAG and RUVBL1-GFP and data from *in vitro* reconstituted complex obtained by SPR. (A) Schematic of the PCR genotyping for the cassette integrated at the RPAP3 genomic locus in HCT116 cells that already express RUVBL1-GFP. The PCR products are indicated in red below the cassette (1 and 2). The gel image shows the results of PCR genotyping. The size of the DNA ladder is indicated on the left. (B) Western blot with anti-RPAP3 and anti-GAPDH antibodies on extracts from parental and HCT116 RPAP3-dTAG cells treated or not with dTAG-V1 for 2h. The size of the protein ladder is indicated on the left. The names of the detected proteins are on the right. (C) Volcano plot representing the enrichment in the RUVBL1-GFP IP versus the control IP (x axis; Log_2_(ratio)) as a function of the p-value (y axis; -Log10), in HCT116 RPAP3-dTAG RUVBL1-GFP cells in untreated condition. Each dot represents a protein that is colored according to the legend above the panel D. (D) Volcano plot representing the enrichment of the mRNAs found in the RUVBL1-GFP IP over the control IP (x axis, Log_2_ (ratio)) as a function of p-values (-Log10, y axis), in HCT116 RPAP3-dTAG RUVBL1-GFP cells in untreated condition. Each dot represents a mRNA and is colored according to the legend above the graph. (E) Affinity (KDss) and Kinetic (*k_a_*, *k_d_*, K_D_) interaction parameters as determined by SPR in Figure 8.

